# Endothelial Cu Uptake Transporter CTR1 Senses Disturbed Flow to Promote Atherosclerosis through Cuproptosis

**DOI:** 10.1101/2025.01.27.634587

**Authors:** Varadarajan Sudhahar, Zhen Xiao, Archita Das, Dipankar Ash, Shikha Yadav, Carson D. Matier, Aidan T. Pezacki, Barun Chatterjee, Olga A. Antipova, Stefan Vogt, Malgorzata McMenamin, Stephanie Kelley, Gabor Csanyi, Jaekwon Lee, Hanjoong Jo, Christopher J. Chang, Jianghong Rao, Jack H. Kaplan, Masuko Ushio-Fukai, Tohru Fukai

## Abstract

Endothelial cells (ECs) lining blood vessels sense disturbed blood flow (D-flow), which drives mitochondrial dysfunction and atherosclerosis. Copper (Cu) is an essential micronutrient, and its disruption of homeostasis has been implicated in atherosclerosis. Cellular Cu levels are tightly controlled by Cu transport proteins including the Cu importer CTR1. Cuproptosis is a recently discovered form of regulated cell death triggered by mitochondrial Cu accumulation, but its endogenous stimulants and role in atherosclerosis remain unknown. Using EC-specific CTR1-deficient mice and cultured ECs, we show that endothelial CTR1 responds to D-flow by increasing mitochondrial Cu levels through its interaction with the mitochondrial Cu transporter SLC25A3 at caveolae/lipid rafts. This leads to the aggregation of lipoylated mitochondrial proteins, mitochondrial dysfunction, and cuproptosis, thereby exacerbating atherosclerosis. Importantly, mitochondria-targeted Cu-chelating nanoparticles effectively mitigate D-flow-induced cuproptosis and atherosclerosis, highlighting the endothelial CTR1-SLC25A3-mitochondrial Cu axis as a potential therapeutic target.

---

Endothelial cells (ECs) are subjected to mechanical forces from blood flow, which elicit distinct cellular responses depending on the vascular region. Laminar flow (L-flow), prevalent in straight vascular segments, limits EC activation and provides protection against atherosclerosis. Conversely, disturbed flow (D-flow), which occurs primarily in arterial curves and branch points, enhances EC activation, promoting endothelial dysfunction and atherosclerotic lesion development. D-flow-induced endothelial dysfunction is characterized by increased inflammation, mitochondrial dysfunction, enhanced permeability, and cell death (*1, 2*). Mitochondrial dysfunction, characterized by impaired respiration and disrupted mitochondrial dynamics, plays a critical role in D-flow-mediated atherosclerosis progression (*3, 4*). However, the mechanisms through which ECs sense D-flow to trigger mitochondrial dysfunction and cell death remain poorly understood.

Copper (Cu) is an essential micronutrient and cofactor for various cellular processes, including mitochondrial respiration, antioxidant defense, and cellular signaling (*5*), making it crucial for maintaining vascular health (*6*). However, excess levels of Cu are toxic. Elevated Cu has been observed in human atherosclerotic plaques (*7*) and plays a critical role in inflammation and pathogenesis of atherosclerosis (*6, 8–10*). Both Cu deficiency and excess have been associated with atherosclerosis (*11*). While Cu has been shown to promote vascular inflammation and atherosclerosis (*12*), the use of Cu chelators in murine models has been shown to mitigate these effects (*13, 14*). These observations highlight the importance of maintaining proper Cu homeostasis to prevent the development of atherosclerotic lesions. Recent studies reveal that mitochondrial Cu overload triggers a unique form of programmed cell death called cuproptosis (*15, 16*). Tsvetkov et al. demonstrated that mitochondrial Cu accumulation, induced by the Cu ionophore elesclomol, induces Cu binding to lipoylated components of the TCA cycle (e.g. dihydrolipoamide S-acetyltransferase (DLAT)), resulting in lipoylated protein aggregation, loss of iron (Fe)-sulfur (S) cluster proteins, including electron transport chain enzymes (*17, 18*). These Cu-dependent processes lead to mitochondrial dysfunction, proteotoxic stress, and cell death (*16, 19*). Unlike ferroptosis, which is also implicated in atherosclerosis (*20*) and driven by iron-dependent lipid hydroperoxides without mitochondrial dysfunction (*21*), cuproptosis is specifically characterized by mitochondrial Cu overload. Notably, Cu chelators prevent elesclomol (Cu ionophore)-induced, Cu-mediated cell death, whereas inhibitors of other cell death pathways, such as necroptosis, ferroptosis, and apoptosis, do not prevent elesclomol-induced cell death (*16*). This underscores the distinct nature of cuproptosis compared to other cell death mechanisms. However, the endogenous triggers of cuproptosis and the role of Cu in D-flow-induced endothelial and mitochondrial dysfunction which drive atherosclerosis remain unknown.

Cu homeostasis is tightly regulated to avoid toxicity. Cellular Cu uptake is primarily mediated by the Cu uptake transporter CTR1 (*22*), which supports essential functions such as mitochondrial activity, superoxide dismutase (SOD)1 activity, secretory cuproenzyme activity, and cellular signaling (*5*). CTR1 forms a selective trimeric pore at the plasma membrane, with conserved methionine residues critical for Cu transport (*23, 24*). Our previous work demonstrated that CTR1 is localized at plasma membrane caveolae/lipid rafts in vascular cells (*25*) and that vascular endothelial growth factor (VEGF) stimulates CTR1 internalization to early endosomes, promoting angiogenesis in ECs (*26*). Additionally, mechanical stress, including shear stress, activates various endothelial membrane molecules, such as ion channels (*27*), integrins (*28*), and VEGFR2 (*29*), but whether CTR1 responds to D-flow to regulate endothelial function is unknown. Mitochondrial Cu transport is mediated by the carrier protein SLC25A3, which transports Cu and phosphate into the mitochondrial matrix (*10*). Depletion of SLC25A3 reduces mitochondrial Cu accumulation, impairing cytochrome c oxidase (COX) activity, which is reversible with Cu supplementation (*30, 31*). Mutations in SLC25A3 are linked to mitochondrial diseases, including cardiomyopathy and lactic acidosis (*32*). Although SLC25A3’s role in mitochondrial Cu regulation has been reported, its possible involvement in D-flow-induced mitochondrial dysfunction remains unexplored.

In this study, we utilize inducible EC-specific Ctr1-deficient (*Ctr1*^iECKO^) mice on *Apoe*^−/−^ background (*Ctr1*^iECKO^/*Apoe*^−/−^) and a recently developed mitochondria-targeted Cu-depleting nanoparticle (mitoCDN)(*33*) to demonstrate that endothelial CTR1 responds to D-flow by increasing mitochondrial Cu levels. This process involves SLC25A3 recruitment to plasma membrane caveolae/lipid rafts (C/LR), where it interacts with CTR1, leading to mitochondrial Cu overload, cuproptosis, and accelerated atherosclerosis. Our findings identify mitochondrial Cu as a central mediator of D-flow-induced atherosclerosis and highlight the CTR1/SLC25A3 axis as a promising therapeutic target to mitigate mitochondrial dysfunction, cuproptosis, and atherosclerosis.

## Results

### Cu accumulation induced by D-flow via endothelial CTR1 drives inflammation and atherosclerosis

To examine the effects of hemodynamics on Cu levels *in vivo*, we analyzed the spatial distribution of Cu in aortas from *Apoe*^−/−^ mice fed a high-fat diet (HFD) for one month. Using synchrotron-based X-ray fluorescence microscopy (XFM), we quantified Cu, iron (Fe), and zinc (Zn) levels, as these metals are implicated in atherosclerosis (*6, 34, 35*). XFM analysis revealed significantly elevated Cu levels in the lesser curvature of the aorta, a region exposed to D-flow, compared to the greater curvature of the aorta, a region exposed to the atheroprotective stable laminar flow (L-flow) (Fig. 1A). In contrast, Fe and Zn levels showed no significant differences between these regions. These findings suggest that D-flow promotes Cu accumulation specifically in the atheroprone lesser curvature of the aortic arch, highlighting a potential link between D-flow, Cu accumulation, and atherosclerosis.

**Figure 1.**
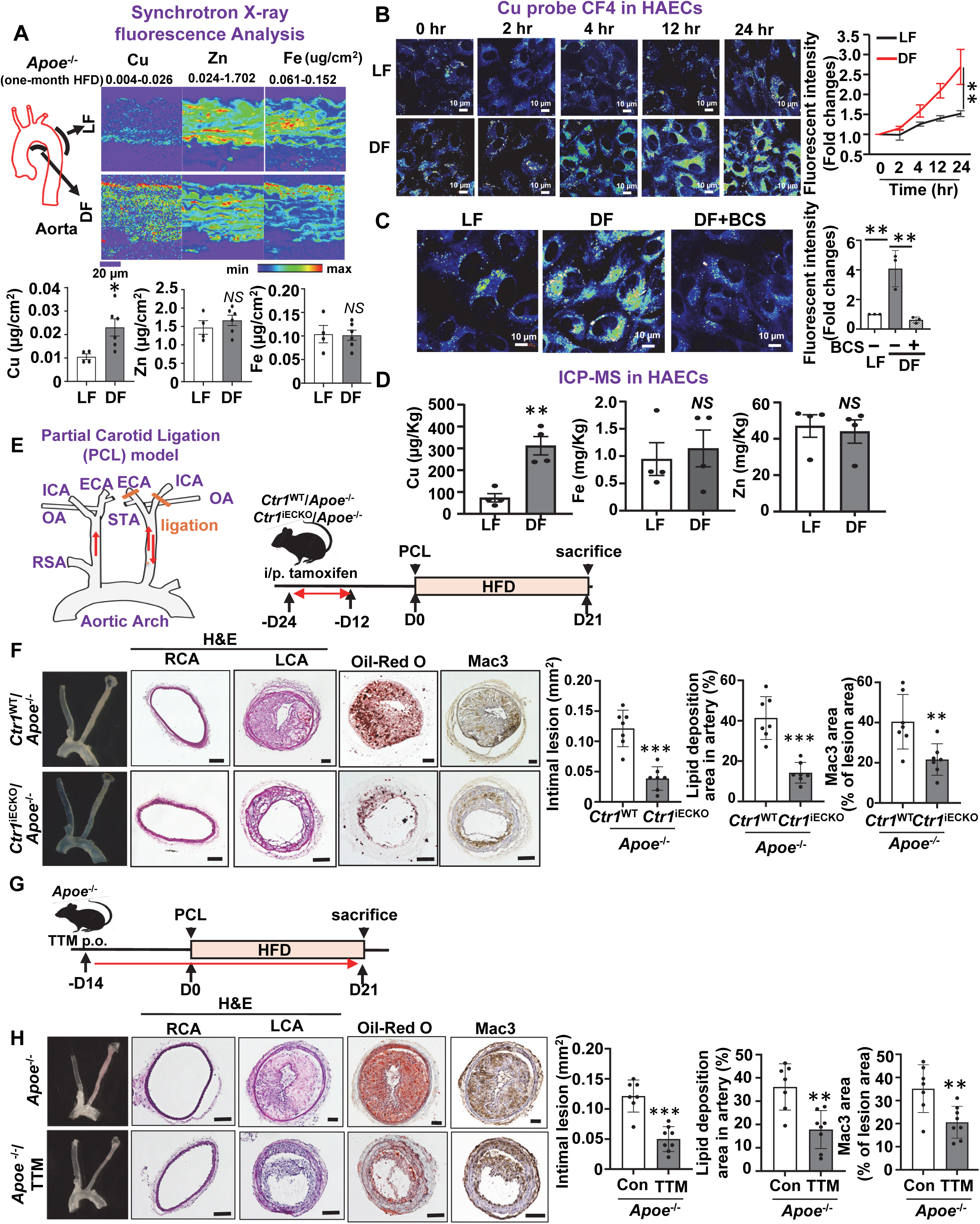
D-flow, but not L-flow, induces Cu accumulation via CTR1 in ECs, which drives inflammation and atherosclerosis. **A.** *Apoe*^−/−^ mice were fed a high fat diet (HFD) for 3 weeks. Synchrotron X-ray fluorescence microscopy (XFM) analysis of the lesser curvature (D-flow, DF) and greater curvature (L-flow, LF) of aorta. The minimal and maximal metal content displayed in micrograms per square centimeter is shown. Scale bars, 20 μm. Quantification of Copper (Cu), Iron (Fe) and Zinc (Zn) accumulation are shown in right (n=4-6). **B-C**. Human aortic endothelial cells (HAECs) were exposed to laminar flow (LF, 15 dyne/cm^2^) or disturbed flow (DF, ± 5 dyne/cm^2^, 1 Hz frequency) conditions for various times (**B**). HAECs were treated with Cu chelator, Bathocuproinedisulfonic acid (BCS, 200 µM for 48 hrs) and exposed to LF and DF for 24 hrs (**C**). Cells incubated with the Cu-specific fluorescent sensor CF4 (2 µM) for 5 min at 37°C were imaged by confocal microscopy (n=3). **D.** ICP-MS measurement for Cu, Fe and Zn contents in HAEC lysates after DF or LF for 24 hrs (n=4). **E.** Partial carotid ligation (PCL) surgery on left carotid artery (LCA) and schematic diagram of experimental design for (**F**). Male and female *Ctr1*^WT^/*Apoe*^−/−^ and *Ctr1*^iECKO^/*Apoe*^−/−^ mice were treated with tamoxifen, followed by PCL surgery on LCA and high fat diet (HFD) for 21 days. **F.** Representative images of cross sections of right carotid artery (RCA) or LCA stained with hematoxylin and eosin (H&E), Oil Red O, or Mac-3 (n=6-7) Scale bar: 100 μm. Right panel: Quantification of lesion size, lipid deposition, and macrophage infiltration. **G.** Schematic diagram of experimental design for (**H**). *Apoe*^−/−^ mice were treated with Cu chelator tetrathiomolybdate (TTM) for 14 days, followed by PCL surgery on LCA and HFD for 21 days. **H.** Representative images of cross sections of RCA or LCA stained with H&E, Oil Red O, or Mac-3 (n=6-7) Scale bar: 100 μm. Right panel: Quantification of lesion size, lipid deposition, and macrophage infiltration.

*In vitro*, we measured intracellular Cu levels ([Cu]i) in human aortic endothelial cells (HAECs) exposed to D-flow (±5 dyne/cm²) or L-flow (15 dyne/cm²) using the well-established *in vitro* shear stress models, using the cone-and-plate and parallel-plate flow chambers (*36*). D-flow induced a significant time-dependent increase in [Cu]i levels, as detected by the Cu-specific fluorescent sensor Cu fluor-4 (CF4)(*37*), compared to L-flow (Fig. 1B). This D-flow-induced Cu elevation was abolished by pre-treatment with the extracellular Cu chelator bathocuproine disulfonate (BCS) (Fig. 1C). Furthermore, D-flow increased CTR1 protein levels without altering its mRNA expression (Figs. S1A, S1B). Consistent with these findings, inductively coupled plasma mass spectrometry (ICP-MS) revealed elevated Cu levels in HAECs exposed to D-flow, whereas Fe and Zn levels remained unchanged (Fig. 1D).

Since CTR1 is a major Cu uptake transporter localized at the plasma membrane, we examined the role of CTR1 in D-flow-mediated atherosclerosis *in vivo* using a partial carotid artery ligation (PCL) model in *Apoe*^−/−^ mice fed a HFD for 21 days (*36*). PCL surgery was performed on the left carotid artery (LCA) (Fig. 1E). Immunofluorescence (IF) staining demonstrated a significant upregulation of CTR1 in CD31-positive ECs in the ligated LCA compared to the non-ligated right carotid artery (RCA) (Fig. S2). To directly assess the role of endothelial CTR1, we generated inducible EC-specific *Ctr1*-deficient mice on an *Apoe*^−/−^ background (*Ctr1*^iECKO^/*Apoe*^−/−^) by crossing *Ctr1*^fl/fl^*/Apoe*^−/−^ (*38*) with *Cdh5*-*CreER^T2^*/*Apoe*^−/−^ (*39*)(Fig. S3A). Selective deletion of Ctr1 in ECs was verified via qPCR (Fig. S3B).

*Ctr1*^WT^ (*Ctr1*^fl/fl^, or *Cdh5*-*CreER^T2^*)/*Apoe*^−/−^ and *Ctr1*^iECKO^/*Apoe*^−/−^ were fed on a high fat diet (HFD) for 3 weeks. A partial ligation surgery of the left carotid artery was performed to introduce D-flow in the left common carotid artery (Fig. 1E). At 3-week post-operation, the *Ctr1*^iECKO^ *Apoe*^−/−^ mice exhibited significantly reduced atherosclerotic lesions, as evidenced by Oil Red O staining, and lower macrophage infiltration (Mac3+) compared to WT mice (Fig. 1F). Importantly, this reduction occurred without altering serum lipid profiles (Fig. S4A). Similarly, treatment of *Apoe*^−/−^ mice with the cell-permeable Cu chelator tetrathiomolybdate (TTM) significantly reduced atherosclerosis and inflammation (Fig. 1G, 1H). TTM efficacy was confirmed by decreased activity of the Cu-dependent enzyme ceruloplasmin in serum, a marker of bioavailable Cu (Fig. S5). Serum cholesterol and triglyceride levels remained unaffected by TTM treatment (Fig. S4B). These findings suggest endothelial CTR1 as a key mediator of D-flow-induced Cu accumulation, promoting atherosclerosis and inflammation in a Cu-dependent manner.

### D-flow induces mitochondrial dysfunction and cuproptosis in ECs through CTR1- and Cu-dependent mechanisms

Mitochondrial dysfunction in ECs is a well-established contributor to atherosclerosis (*40*). To evaluate the impact of D-flow on mitochondrial function, we assessed mitochondrial respiration in atherosclerosis-prone regions of the aorta exposed to D-flow, where Cu levels are elevated (Fig. 1A). Using a Seahorse XF24 extracellular flux analyzer, we observed a significant decrease in oxygen consumption rate (OCR) in regions exposed to D-flow compared to those protected by L-flow, in an endothelium-dependent manner (Fig. 2A). Similarly, HAECs exposed to D-flow showed reduced OCR compared to L-flow-treated cells (Fig. 2B). Importantly, this D-flow-induced decrease in OCR was reversed by pre-treatment with TTM or by CTR1 knockdown using siRNA (Fig. 2B).

**Figure 2.**
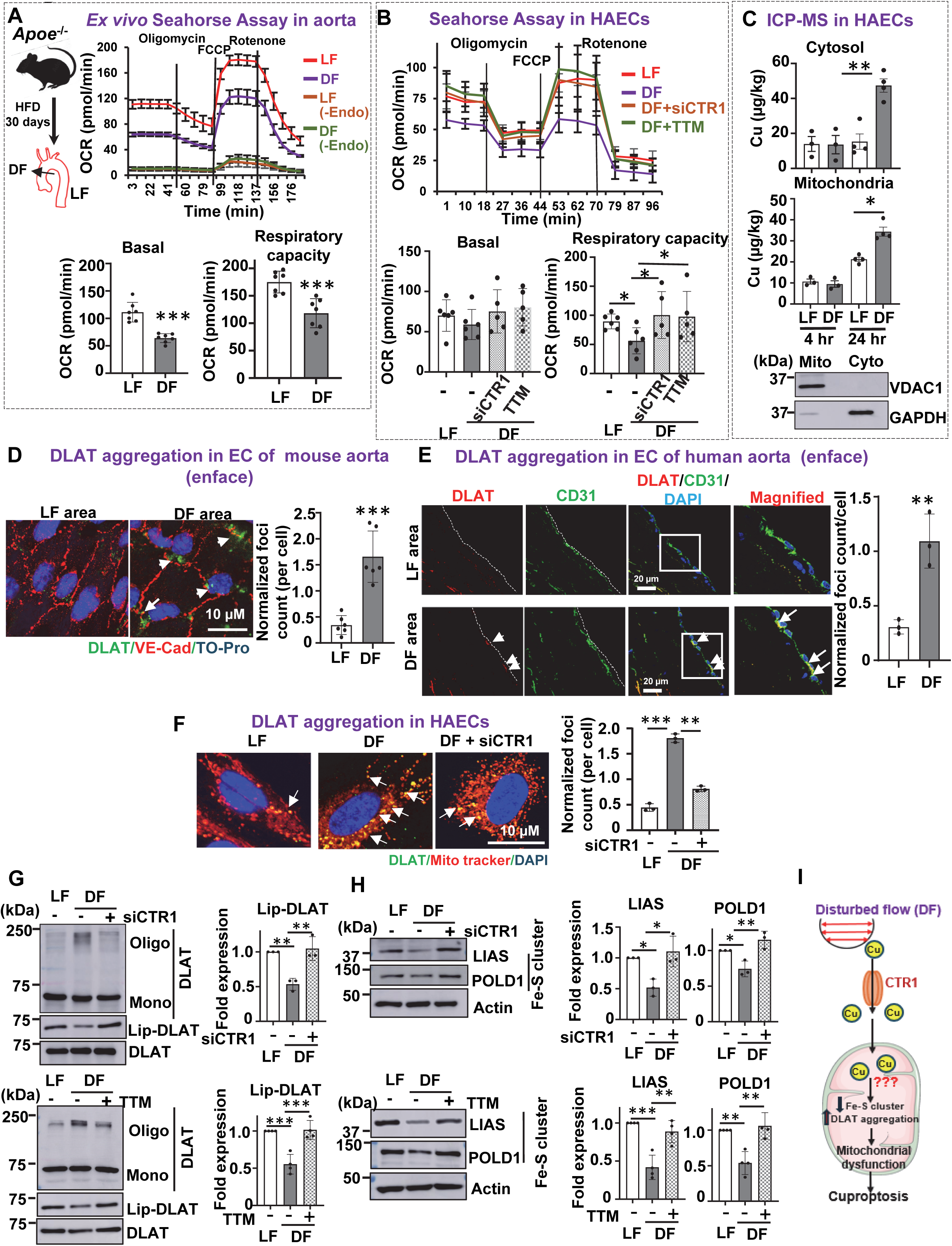
D-flow promotes mitochondrial dysfunction and cuproptosis via CTR1- and Cu-dependent mechanisms in ECs. **A.** *Ex vivo* Seahorse assay to measure mitochondrial O_2_ consumption rate (OCR) in aortic arch (D-flow, DF) and thoracic aorta (L-flow, LF) with and without endothelial removal by denudation, isolated from *Apoe*^−/−^ mice fed with HFD for 30 days (n=7). **B.** OCR in HAECs transfected with control siRNA or CTR1 siRNA, or pretreated with or without Cu chelator, TTM (20nM, 24 hrs) exposed to LF or DF for 48 hrs. (n=5-6). **C.** HAECs exposed to DF or LF for 4 hr or 24 hrs were used to measure Cu content in cytosolic and mitochondrial fraction using ICP-MS (n=3-4). **D.** Left: Representative images of *en face* immunofluorescence staining of DLAT aggregation (green). The endothelium was visualized by VE-Cadherin (VE-Cad)(Red) staining, and nuclei were counterstained with TO-Pro (blue). Right: Quantification of DLAT aggregation in the endothelial layer of aorta exposed to LF or DF (n=5). **E.** Left: Representative images of *en face* immunofluorescence staining with DLAT aggregation (red) in the endothelium of greater curvature (LF) and lesser curvature (DF) of aortic arch of human atherosclerotic aorta. The endothelium was visualized by CD31 (green) and nuclei were counterstained with DAPI (blue) (n=3). Right: Quantification of DLAT aggregation. Details of donor tissues are described in previous reports (*41*). Additionally, patient #3’s information is: 69/F/Caucasian/female, Respiratory failure (cause of death), and Coronary Artery disease/hypertension/kidney disease (Medial history). **F.** HAECs transfected with CTR1 siRNA or control siRNA were exposed to DF for 48 hrs. Representative images of *en face* immunofluorescence staining DLAT aggregation (green, DLAT; red, Mitotracker; blue, DAPI). **G and H.** HAECs were transfected with CTR1 siRNA or control siRNA or pretreated with 20 nM TTM for 24 hrs and exposed to DF for 48 hrs. Cell lysates were immunoblotted with the indicated antibodies (n=3-4). **I.** Schematic model showing that DF promotes Cu accumulation in cytosol and mitochondria, as well as mitochondrial dysfunction and cuproptosis in a CTR1/Cu-dependent manner in ECs.

To further investigate the role of Cu in mitochondrial dysfunction, ICP-MS analysis of subcellular fractions revealed a significant increase in Cu levels in both the cytosol and mitochondria under D-flow conditions, while Fe and Zn levels remained unchanged (Fig. 2C and Fig. S6). Under L-flow, a moderate increase in mitochondrial Cu levels was observed at 24 hours without significant changes in cytosolic Cu levels. This moderate increase may affect mitochondrial Cu-dependent enzymes, such as COX (Fig. 2C) without causing cuproptosis (see below). Mitochondrial fractionation was validated by the presence of VDAC1 in mitochondrial fractions and its absence in cytosolic fractions (Fig 2C).

We next investigated whether D-flow induces cuproptosis, a recently identified form of mitochondrial Cu-mediated cell death. This process involves the accumulation of mitochondrial Cu, which binds directly to lipoylated components of the TCA cycle, such as DLAT, leading to lipoylated protein aggregation, loss of iron-sulfur cluster proteins, and subsequent cell death (*16*). Thus, we examined whether DLAT aggregation is observed in atherosclerotic aortas from mice and humans, as well as in cultured ECs exposed to D-flow. Immunofluorescence staining revealed that D-flow promotes DLAT aggregation foci in the EC layer of isolated mouse aortas, as shown by *en face* staining (Fig. 2D). Analysis of the lesser curvature of human atherosclerotic aortas, a region exposed to D-flow, demonstrated increased DLAT aggregation in ECs compared to regions exposed to L-flow (Fig. 2E). Details of human donor tissues have been previously described (*41*). Similarly, D-flow-induced DLAT aggregation was observed in HAECs, as evidenced by the co-localization of DLAT with MitoTracker (Fig. 2F). This aggregation was significantly reduced by CTR1 knockdown (Fig. 2F) or TTM treatment (data not shown), suggesting a Cu-dependent process.

We further confirmed D-flow effects on cuproptosis-related responses, demonstrating that D-flow increases DLAT oligomerization under non-reducing conditions, accompanied by decreased DLAT lipoylation (Fig. 2G). The knockdown of CTR1 or TTM treatment mitigated D-flow-induced DLAT oligomerization, supporting a Cu-dependent mechanism. Additionally, D-flow reduced the expression of Fe-S cluster proteins (LIAS, POLD1, and Complex I), which was restored by Cu chelation or CTR1 knockdown (Fig. 2H). Reduced lipoic acid synthetase (LIAS) expression may account for some decreased DLAT lipoylation, as LIAS is a key regulator of this process (*42*). D-flow also downregulated oxidative phosphorylation complex proteins (Complexes I, III, and IV), which was prevented by CTR1 siRNA or TTM treatment (Figs. S7A, S7B).

D-flow-induced cell death was then evaluated in cultured ECs. D-flow increased cell death, as evidenced by the CCK8 assay for cytotoxicity and LDH release in HAECs. These effects were significantly attenuated by TTM treatment or CTR1 knockdown (Fig. S8). Inhibitors that target alternative cell death pathways, including ferroptosis (Fer-1) and necroptosis (Nec-1), failed to prevent D-flow-induced cell death, whereas the pan-caspase inhibitor Z-VAD-FMK partially rescued viability (Fig. S8). These findings suggest that while apoptosis is partially involved, cuproptosis is the predominant pathway driving D-flow-induced EC death. These results demonstrate that D-flow promotes mitochondrial Cu accumulation, dysfunction, and cuproptosis in ECs.

### Mitochondrial Cu elevation induced by D-flow contributes to the promotion of cuproptosis and atherosclerosis

To explore the impact of mitochondrial Cu in D-flow-mediated mitochondrial dysfunction, cuproptosis, and atherosclerosis, we employed a mitochondria-targeted Cu-depleting nanoparticle (mitoCDN) (Fig. 3A) (*33*). MitoCDN selectively reduces mitochondrial Cu levels without altering systemic Cu concentrations (*33*) and comprises two main components: a Cu-depleting moiety (CDM) and a semiconducting polymer nanoparticle (SNP) (Fig. 3A). In cultured ECs, treatment with mitoCDN partially reversed D-flow-induced reductions in OCR and reduced endothelial cell death (Figs. 3B, 3C, and 3D). Furthermore, mitoCDN reversed D-flow-induced cuproptosis-related changes, including DLAT aggregation, decreased DLAT lipoylation, and loss of Fe-S cluster and oxidative phosphorylation complex proteins, to levels comparable to those observed under L-flow conditions (Figs. 3E, 3F, and S7C). *In vivo*, mitoCDN treatment inhibited D-flow-induced cuproptosis markers, including DLAT aggregation and mitochondrial dysfunction, as evidenced by *ex vivo* OCR measurements in aorta (Figs. S9A, 3G, and S9B). In a PCL model, intravenous mitoCDN administration (Fig 3H) reduced D-flow-induced cell death (Figs. 3H, 3I, 3J), vascular inflammation, and atherosclerotic lesion formation in *Apoe*^−/−^ mice (Figs. 3H, 3K, and 3L) without altering serum cholesterol and triglyceride levels (Fig. S4C). Collectively, these findings suggest that D-flow-induced mitochondrial Cu accumulation promotes mitochondrial dysfunction and cuproptosis, contributing to the development of atherosclerosis.

**Figure 3.**
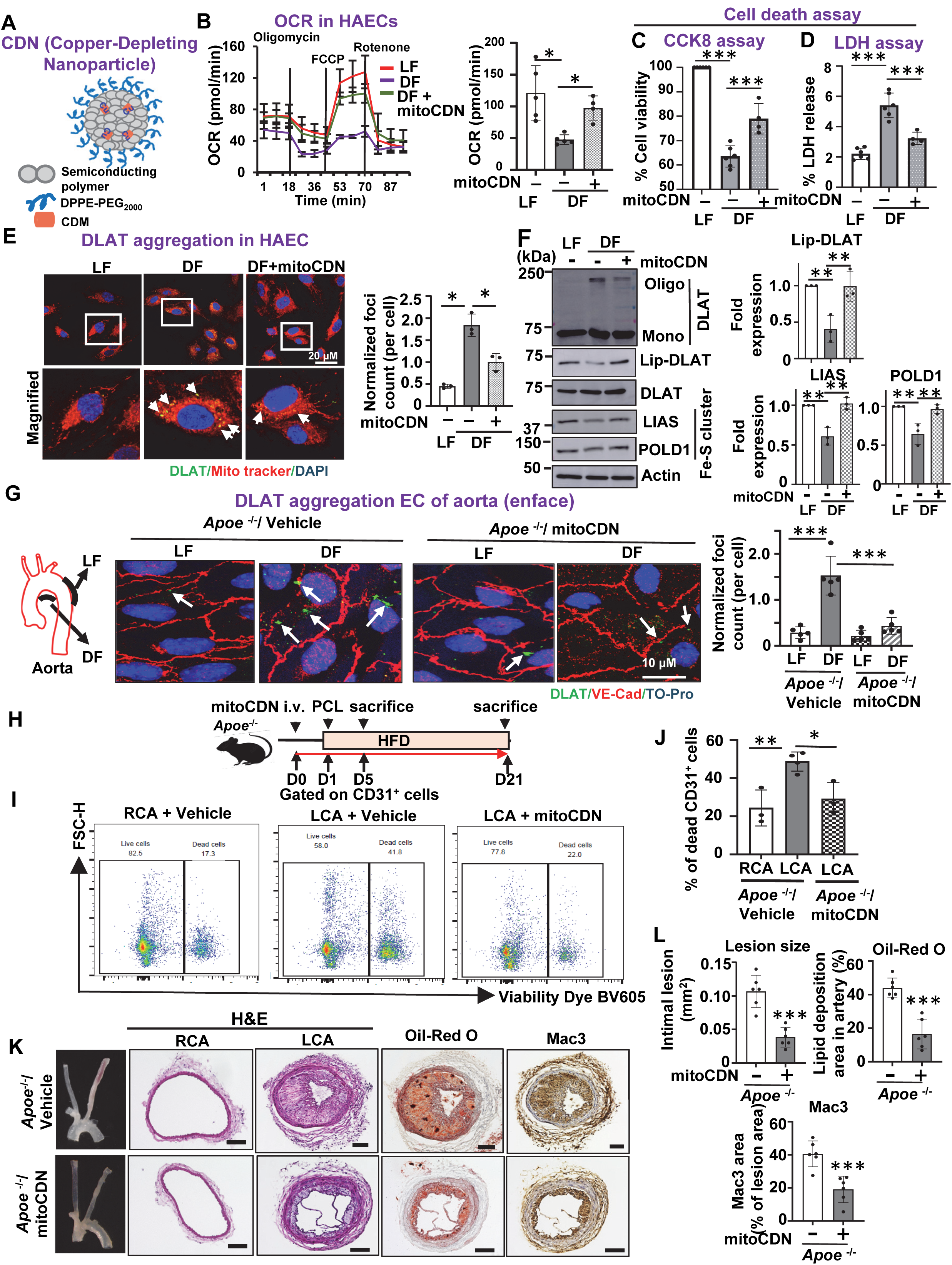
D-flow-induced mitochondrial Cu elevation contributes to cuproptosis and atherosclerosis. **A.** Molecular components of mitoCDN and illustrated nanoparticle formulation. **B-F.** HAEC were pretreated with 1 µM mitoCDN for 1 hr and exposed to DF (D-flow) for 48 hrs. Mitochondrial O_2_ consumption rate (OCR) measured by a Seahorse analyzer (n=5-6)(**B**). Cell death was measured by CCK8 assay (n=5) (**C**) or LDH release assay (n=4)(**D**). Confocal immunofluorescence imaging of DLAT aggregation (green, DLAT; red, Mitotracker; blue, DAPI)(n=3)(**E**). Cell lysates were immunoblotted with the indicated antibodies (n=3)(**F**). **G.** *Apoe*^−/−^ mice were treated with mitoCDN (1.35mg/kg) every 3 days with a total of 10 doses and fed with HFD for 30 days. Representative images of en face staining for DLAT aggregation (green), VE-Cad (endothelium, red) and DAPI (nucleus, blue) in the endothelial layer with LF (L-flow) or DF. (n=5). **H.** Schematic diagram of experimental design for **I** to **L**. DF was induced in the LCA of *Apoe*^−/−^ mice using PCL surgery, while the contralateral RCA was used as an internal control. Ligated mice were fed with HFD and treated with mitoCDN (i.v.) (one day prior to surgery and every three days after PCL). **I** and **J.** LCA or RCA from 3 mice were pooled and digested at 4 days after PCL. CD31^+^ ECs were isolated and used to analyze cell viability using flow cytometry with viability dye, BV605. Right: Quantification of % dead ECs. (n=3). **K** and **L.** Representative images of cross sections of RCA or LCA stained with H&E, Oil Red O, and Mac-3 at 3 weeks after PCL (n=6) Scale bar: 100 μm. Quantification of lesion size, lipid deposition, and macrophage infiltration (Right) (n=6).

### D-flow facilitates the translocation of SLC25A3 to lipid rafts, its interaction with CTR1, and promotes mitochondrial Cu accumulation, leading to cuproptosis in ECs

To investigate how D-flow increases mitochondrial Cu levels through CTR1, we examined the role of the mitochondrial Cu transporter SLC25A3 (*10*). Previous studies have shown that D-flow elevates cholesterol levels in plasma membrane caveolin1 (Cav)1-enriched caveolae/lipid rafts (C/LR), recruiting mitochondrial ATP synthase (*43*), and that CTR1 is also found within these domains (*25*). We therefore examined whether D-flow recruits the mitochondrial protein SLC25A3 to the C/LR. Using detergent-free sucrose density gradient centrifugation (*25, 44*), we isolated C/LR fractions from ECs exposed to either D-flow or L-flow. Western blot analysis showed that under L-flow conditions, CTR1 was localized in both C/LR and non-C/LR fractions, whereas SLC25A3 was restricted to non-C/LR fractions (Fig.4A). However, after 24 hours of D-flow exposure, SLC25A3, along with ATP5A1 (a component of ATP synthase), was translocated to the C/LR fractions, while CTR1 levels in these fractions remained unchanged (Figs. 4B and 4C).

**Figure 4.**
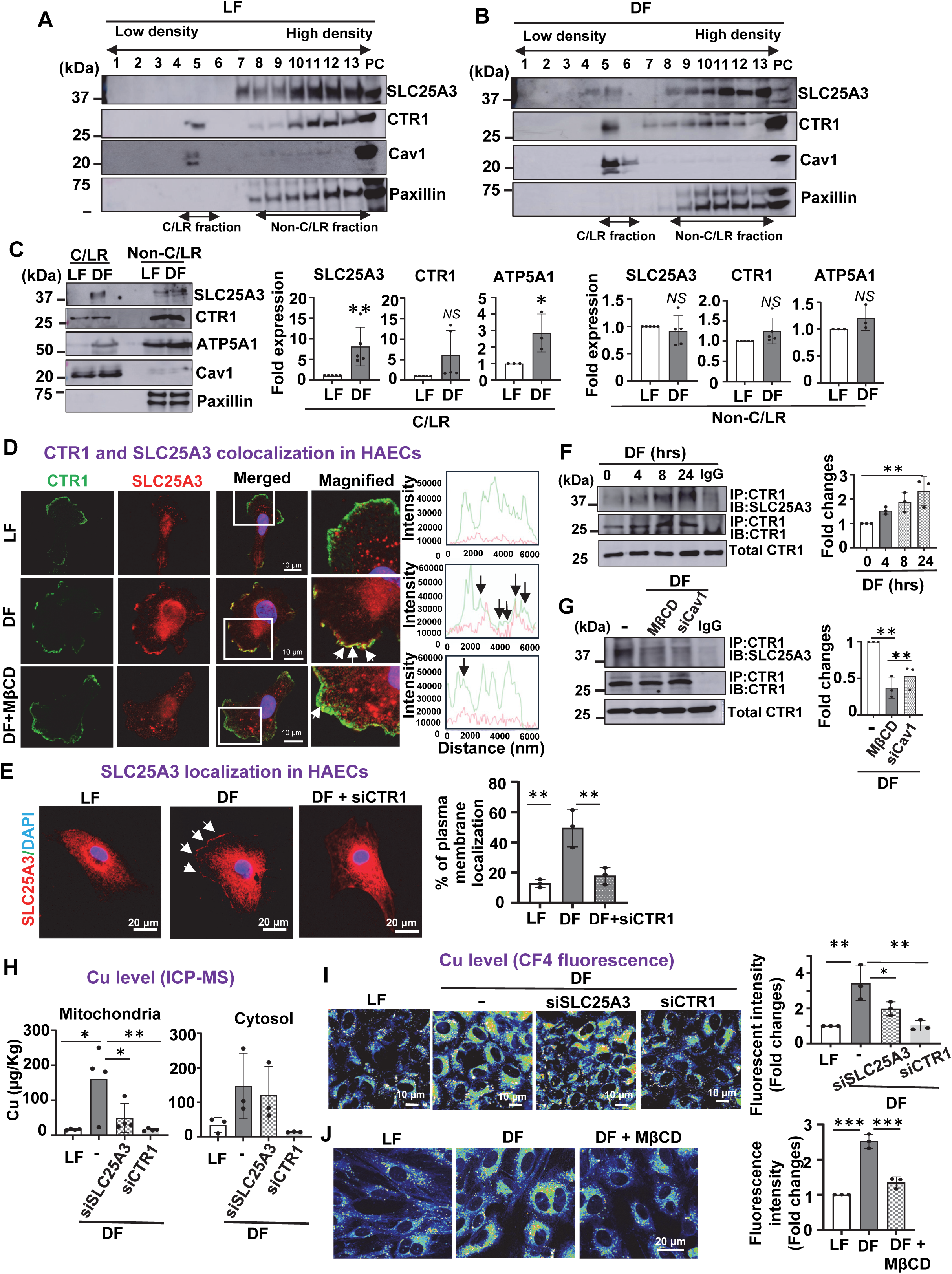
D-flow promotes SLC25A3 translocation to lipid rafts, interaction with CTR1, and mitochondrial Cu accumulation, leading to cuproptosis in ECs. **A** and **B**. HAECs exposed to LF (L-flow) or DF (D-flow) for 24 hrs were used to isolate caveolae/lipid raft (C/LR) using sucrose gradient centrifugation. Fractions from the top (fraction 1) to the bottom (fraction 13) were immunoblotted with antibodies as indicated. **C.** Equal amounts of C/LR (fraction 5) or non-C/LR (fraction10-13) were immunoblotted with antibodies as indicated (n=5). ATP5A1, a component of ATP synthase, is used as a marker of the mitochondrial inner membrane. **D.** HAECs transfected with CTR1-flag plasmid were pretreated with or without Methyl-β-cyclodextrin (MβCD, 10 mM, 1 hr) and then exposed to DF for 24 hrs. Cells were immunostained for Flag (green), SLC25A3 (red) and DAPI (blue) (n=3). **E.** HAECs transfected with control or CTR1 siRNAs were exposed with DF for 24 hrs. Cells were immunostained with SLC25A3 (green) and DAPI (blue). Right: Quantification for % plasma membrane localization of SLC25A3 (n=3). **F** and **G**. HAECs were exposed to DF for indicated time (**F**) or transfected with control or caveolin-1 (Cav1) siRNAs with or without pretreated with MβCD (10 mM, 1 hr) (**G**). These ECs were exposed to DF for 24 hrs. Lysates were immunoprecipitated with CTR1 antibody or IgG, followed by immunoblotted with SLC25A3 or CTR1 antibody. **H, I,** and **J.** HAECs transfected with control or SLC25A3 or CTR1 siRNAs exposed to DF for 24hrs were used to measure Cu contents in cytosol and mitochondrial fractions using ICP-MS (**H**) or [Cu]i levels using CF4 fluorescence probe (**I**). (n=3-4). HAECs treated with or without MβCD (10 mM, 1 hr) exposed to DF for 24 hrs were used to measure [Cu]i levels using CF4 fluorescence probe (**J**)(n=3).

Immunofluorescence analysis revealed that in HAECs exposed to L-flow for 24 hours, CTR1 was localized to the plasma membrane, while SLC25A3 was primarily found near the perinuclear region (Fig. 4D). Under D-flow conditions, SLC25A3 translocated to the plasma membrane, where it colocalized with CTR1. This translocation was inhibited by pretreatment with the cholesterol chelator methyl-β-cyclodextrin (MβCD, 10 mM), which disrupts C/LR structures (Fig. 4D), as well as by CTR1 siRNA (Fig. 4E). Supporting these findings, immunoprecipitation assays further demonstrated that D-flow promoted the interaction between SLC25A3 and CTR1 in a time-dependent manner (Fig. 4F). Notably, plasma membrane C/LR has been shown to structurally and functionally interact with mitochondria via Cav1, regulating cell metabolism (*45, 46*). Importantly, the D-flow-induced interaction between SLC25A3 and CTR1 was disrupted by MβCD or Cav1 siRNA (Fig. 4G). These findings suggest that Cav1 is involved in mediating CTR1-SLC25A3 binding at the mitochondria-C/LR interface (*45, 47*).

To evaluate the functional relevance of the interaction between CTR1 and SLC25A3 in mitochondrial Cu accumulation, we performed mitochondrial fractionation and analyzed Cu levels using ICP-MS. Both CTR1 and SLC25A3 siRNAs inhibited the D-flow-induced increase in mitochondrial Cu levels, while CTR1 siRNA, but not SLC25A3 siRNA, reduced Cu levels in the cytosol fraction (Fig. 4H). In contrast, Fe and Zn levels were unaffected by either CTR1 or SLC25A3 siRNA treatment (Fig. S10). Furthermore, Cu-sensitive fluorescence imaging, which detects intracellular [Cu]i levels in both the cytosol and mitochondria, showed that CTR1 siRNA completely blocked D-flow-induced [Cu]i elevation, while SLC25A3 siRNA only partially inhibited it in ECs (Fig. 4I). Additionally, disruption of C/LR using MβCD significantly reduced [Cu]i elevation induced by D-flow (Fig. 4J). These findings suggest that the CTR1-SLC25A3 axis within the C/LR domain plays a crucial role in mitochondrial Cu accumulation in ECs exposed to D-flow.

Finally, we explored the role of SLC25A3 and C/LR in D-flow-induced cuproptosis in ECs. Knockdown of SLC25A3 or Cav1, as well as MβCD treatment significantly inhibited D-flow-induced DLAT aggregation while restoring DLAT lipoylation and preventing the loss of Fe-S clusters (Fig. 5A, 5B, and 5C). These interventions also mitigated D-flow-induced reductions in oxidative phosphorylation (OXPHOS) complex proteins (Figs. S11A and S11B). Collectively, these findings indicate that D-flow drives mitochondrial Cu accumulation through CTR1-SLC25A3 interactions at the mitochondria-C/LR interface, leading to mitochondrial dysfunction and cuproptosis in ECs. This mechanism likely contributes to the development of flow-dependent atherosclerosis (Fig. 5D).

**Figure 5.**
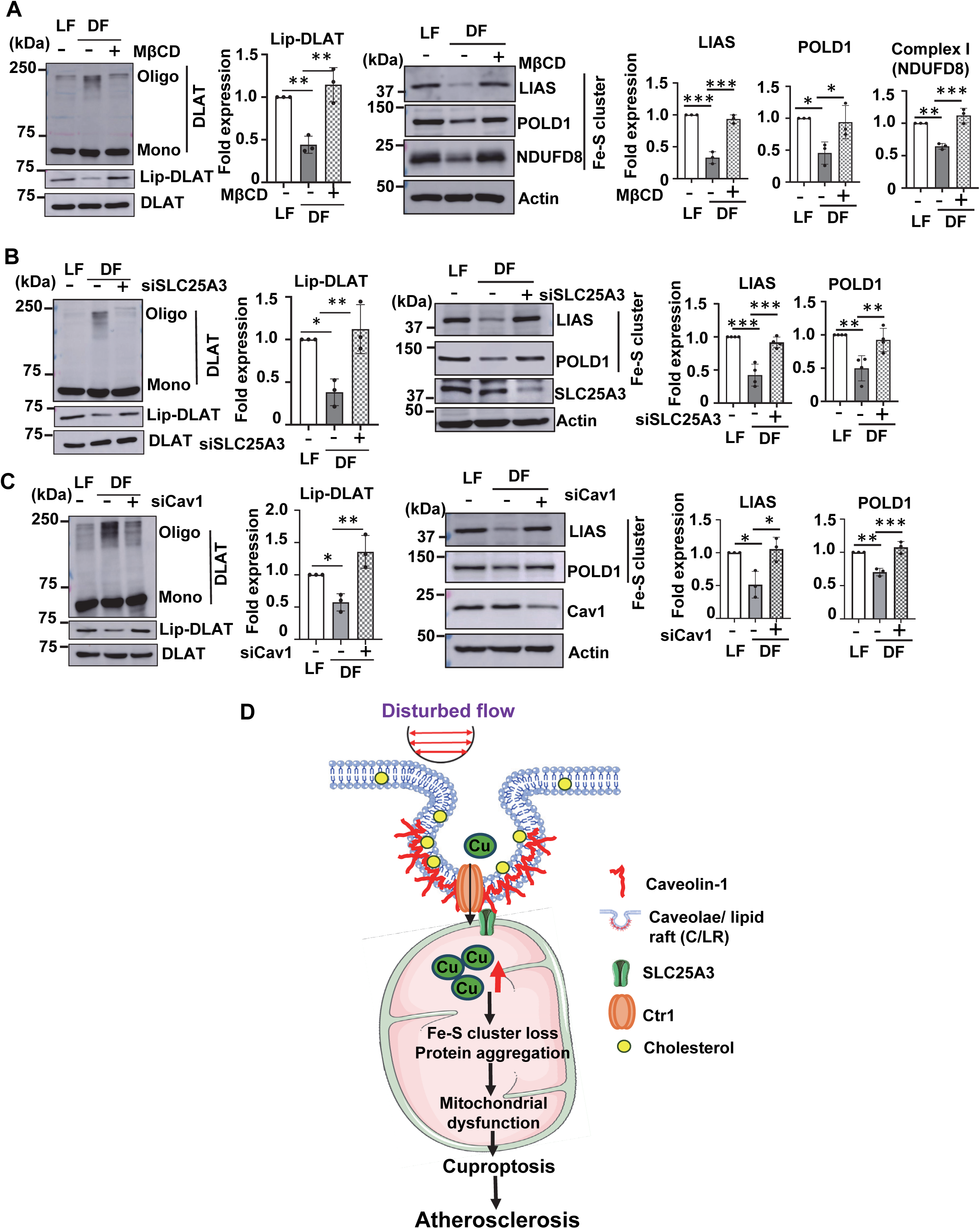
D-flow-induced cuproptosis is mediated through CTR1-SLC25A3- and lipid rafts-dependent mechanisms. A, B, and **C.** HAECs treated with or without MβCD (10 mM, 1 hr) (**A**) or transfected with control or SLC25A3 siRNA (**B**) or Cav1 siRNA (**C**) were exposed to DF (D-flow) or LF (L-flow) for 48 hrs. These ECs were used to measure DLAT aggregation and lipoylation (Left), and protein expression of Fe-S cluster proteins (Right) (n=3). **D.** Schematic summary of the proposed model. Endothelial CTR1 responds to DF by elevating mitochondrial Cu through forming a complex with the mitochondrial Cu transporter SLC25A3 at lipid rafts. This in turn promotes aggregation of lipoylated mitochondrial proteins, mitochondrial dysfunction, and cuproptosis, thereby driving atherosclerosis. Thus, the endothelial CTR1-SLC25A3-mitochondrial Cu axis is a promising therapeutic target for treatment of Cu-dependent vascular diseases such as atherosclerosis.

## Discussion

Cuproptosis is a newly identified form of regulated cell death triggered by the Cu accumulation, particularly in mitochondria, where it disrupts essential metabolic processes following the application of the Cu ionophore elesclomol (*16*). This study provides direct evidence that the endogenous pathophysiological stimulant, D-flow, drives mitochondrial Cu accumulation and triggers cuproptosis in ECs, contributing to mitochondrial dysfunction and atherosclerosis. We also show that D-flow-induced association between the plasma membrane Cu uptake transporter CTR1 and the mitochondrial Cu transporter SLC25A3 facilitate Cu accumulation in the mitochondria. This process initiates mitochondrial dysfunction, leading to cuproptosis and the progression of atherosclerosis. These findings highlight mitochondrial Cu as a pivotal mediator of D-flow-induced vascular pathology and identify the CTR1-SLC25A3 axis as a promising therapeutic target.

D-flow leads to EC dysfunction and inflammation, playing a critical role in the development of atherosclerotic plaques. Cu imbalances, whether due to excess or deficiency and regulated by Cu transporters such as the Cu importer CTR1, have been linked to atherosclerosis. However, the connection between D-flow-induced atherosclerosis and Cu imbalances is not well understood. Using XFM and ICP-MS, we demonstrate that D-flow selectively increases Cu levels in atherosclerosis-prone regions of the aorta and in cultured ECs, without altering iron or zinc levels. This Cu elevation is associated with increased CTR1 protein expression in both *in vivo* and *in vitro* models, supporting the central role for CTR1 in D-flow-induced Cu homeostasis dysregulation. Importantly, *Ctr1*^iECKO^*Apoe^−/−^* mice exhibited reduced D-flow-induced atherosclerotic lesions and inflammation, without affecting systemic lipid profiles, underscoring the significance of endothelial CTR1 in mediating D-flow-induced atherosclerosis. Moreover, Cu chelation with TTM significantly mitigated D-flow-induced atherosclerosis and inflammation, further emphasizing the critical involvement of Cu in this process.

Recent studies have shown that elevated mitochondrial Cu levels induced by the Cu ionophore elesclomol drive a unique form of regulated cell death, termed cuproptosis (*16*). D-flow-induced mitochondrial dysfunction is a hallmark of endothelial dysfunction and atherosclerosis (*36, 40*). However, the role of cuproptosis in this context had not been comprehensively explored. In this study, we present strong evidence that D-flow induces mitochondrial Cu accumulation, which leads to reduced oxygen consumption rates, aggregation of lipoylated DLAT proteins, and reduction of Fe–S (iron–sulfur) cluster proteins (e.g. LIAS, NDUFS8). These changes result in impaired oxidative phosphorylation, a key feature of mitochondrial dysfunction (*40*). Importantly, these effects are reversed by CTR1 knockdown or treatment with Cu chelators, including the newly developed mitochondria-targeted Cu-depleting nanoparticle (mitoCDN) (*33*), emphasizing the pivotal role of mitochondrial Cu in this process. Notably, there is a difference between the pharmacological effects of the Cu ionophore elesclomol, which directly increases mitochondrial and cytosolic Cu (II) (*16*) and the pathophysiological effects of D-flow, which employs CTR1 to transport Cu (I), elevating cellular and mitochondrial Cu (I) levels (*22*). This study demonstrates that D-flow-induced increases in mitochondrial Cu (I) levels can drive cuproptosis and contribute to atherosclerosis through CTR1-dependent pathways in ECs.

Previous studies have shown that CTR1 is localized within C/LR (*25*). Our finding reveals that D-flow promotes mitochondrial Cu accumulation by facilitating the translocation of the mitochondrial Cu transporter SLC25A3 to the plasma membrane C/LR, where it interacts with CTR1. Immunofluorescence and co-immunoprecipitation analyses confirm that CTR1 binds to SLC25A3 in response to D-flow, forming a pathway for Cu transfer from CTR1 to mitochondria. Silencing either CTR1 or SLC25A3 significantly reduces mitochondrial Cu levels. Of note, depletion of SLC25A3 does not affect the D-flow-induced increased cytosolic Cu levels, underscoring its specific role in mitochondrial Cu transport. The C/LR has been shown to interact with mitochondria through Cav1, regulating cell metabolism (*45, 46*). Interestingly, the interaction between CTR1 and SLC25A3 is reduced by MβCD, which disrupts C/LR, or by Cav1 siRNA, indicating that Cav1 mediates and facilitates CTR1-SLC25A3 binding at the mitochondria-C/LR interface (*45, 47*). These findings are consistent with previous studies showing that D-flow enhances cholesterol levels in C/LR and promotes the translocation of mitochondrial ATP synthase to C/LR via Cav1 (*43, 48*). Thus, these results suggest a functional and structural link between mitochondria and plasma membrane C/LR. The mechanism underlying D-flow-induced mitochondria transport to the plasma membrane may involve kinesin superfamily protein (KIF)-microtubule or actin cytoskeleton pathways (*48, 49*), which warrants further investigation.

Our studies identify cuproptosis, a recently characterized form of programmed cell death driven by mitochondrial Cu overload (*16*), as a key mechanism underlying D-flow-induced endothelial and mitochondrial dysfunction. D-flow-mediated mitochondrial Cu accumulation triggers hallmark features of cuproptosis, including lipoylated DLAT aggregation, loss of Fe-S cluster proteins, and reduced expression of OXPHOS complex proteins (*16*). These effects are mitigated by Cu chelation, or by silencing CTR1 or SLC25A3, confirming the Cu-dependent nature of this cell death pathway. Notably, D-flow-induced cell death is not prevented by inhibitors of other cell death pathways, except for apoptosis, emphasizing the specificity of mitochondrial cell death in this context. Furthermore, treatment with mitoCDN (*33*) restores mitochondrial function, reduces DLAT aggregation, and mitigates atherosclerosis *in vivo*. These findings underscore the pivotal role of mitochondrial Cu accumulation and cuproptosis in the development of D-flow-induced atherosclerosis. Mitochondrial Cu homeostasis has also shown potential as a therapeutic target for various diseases, including cancer, as evidenced by the effective suppression of tumor growth using mitoCDN (*33*), and also for Menkes disease (*50*). Further research is required to explore the role of mitochondrial Cu homeostasis disruption in other diseases and cell types.

In conclusion, our studies reveal that the interaction between the Cu uptake transporter CTR1 and the mitochondrial Cu transporter SLC25A3 at the mitochondrial contact site with the plasma membrane lipid rafts is a critical driver of mitochondrial Cu accumulation. This process leads to mitochondrial dysfunction and cuproptosis, ultimately contributing to the progression of atherosclerosis. These findings underscore the potential of the endothelial CTR1-SLC25A3 mitochondrial Cu axis as a therapeutic target for preventing atherosclerosis and other cuproptosis-related diseases, including cardiovascular disorders, cancer, and neurodegenerative diseases.

## Supplemental Figure Legend

**Figure S1.**
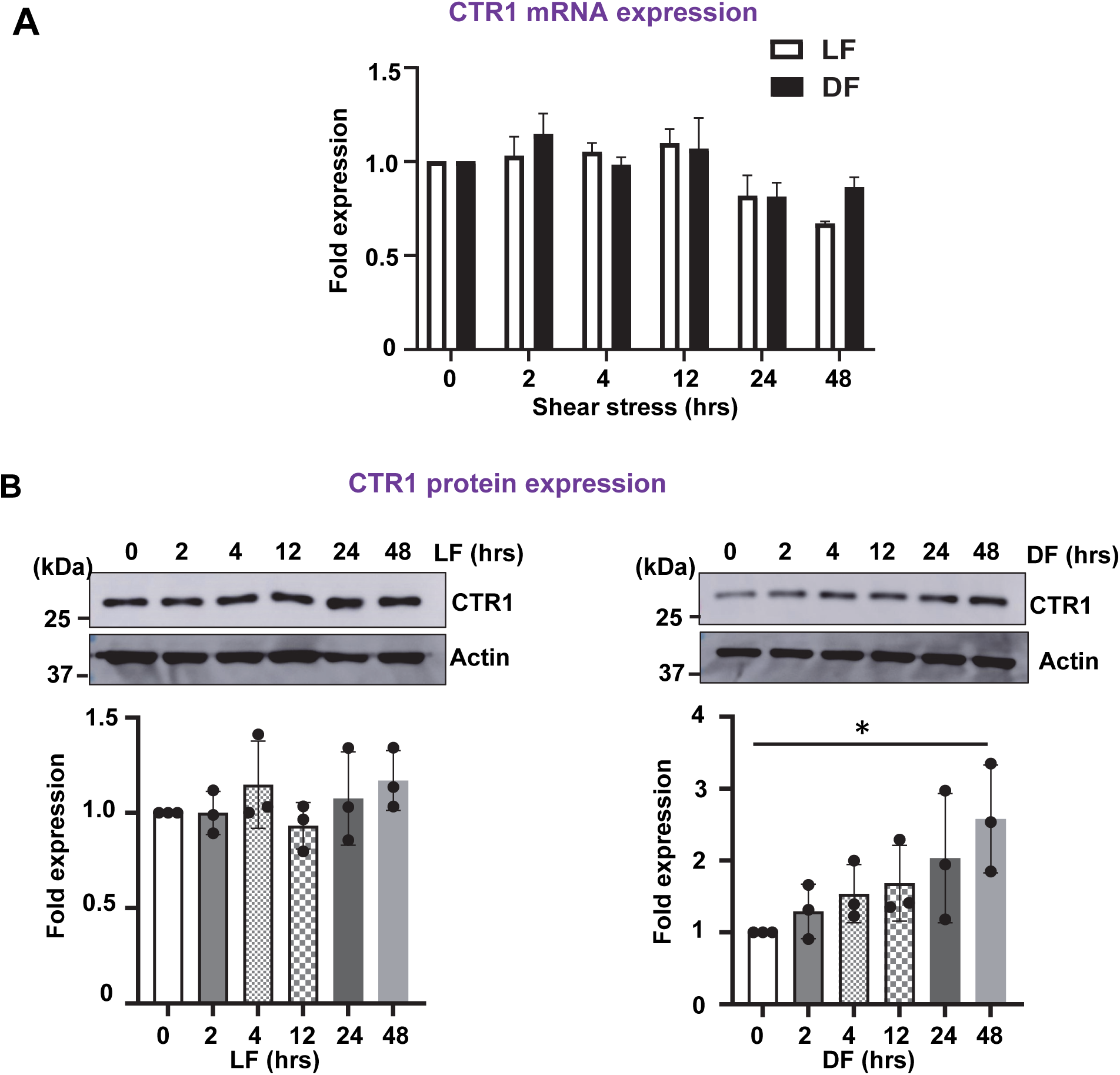
Effect of L-flow and D-flow on Cu uptake transporter CTR1 expression in HAECs. **A** and **B.** HAEC were exposed with LF (L-flow) or DF (D-flow) for indicated times. CTR1 mRNA expressions were measured using quantitative polymerase chain reaction (**A**). To examine CTR1 protein expressions, cell lysates were immunoblotted with the indicated antibodies (n=3) (**B**).

**Figure S2.**
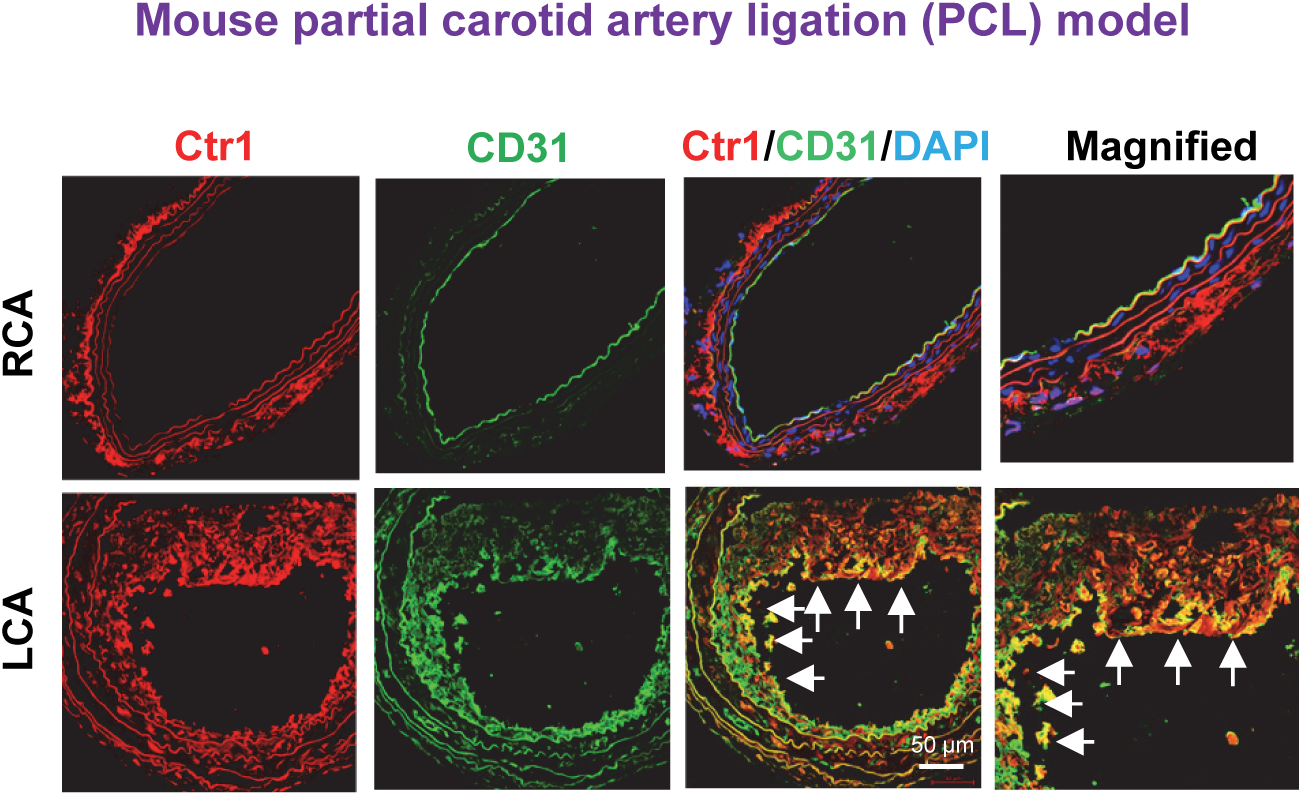
Ctr1 expression is increased in a DF-exposed carotid artery of *Apoe*^−/−^ mice. Immunofluorescence staining of Ctr1 in RCA and LCA of *Apoe*^−/−^ mice 2 weeks after PCL surgery on LCA (n=3).

**Figure S3.**
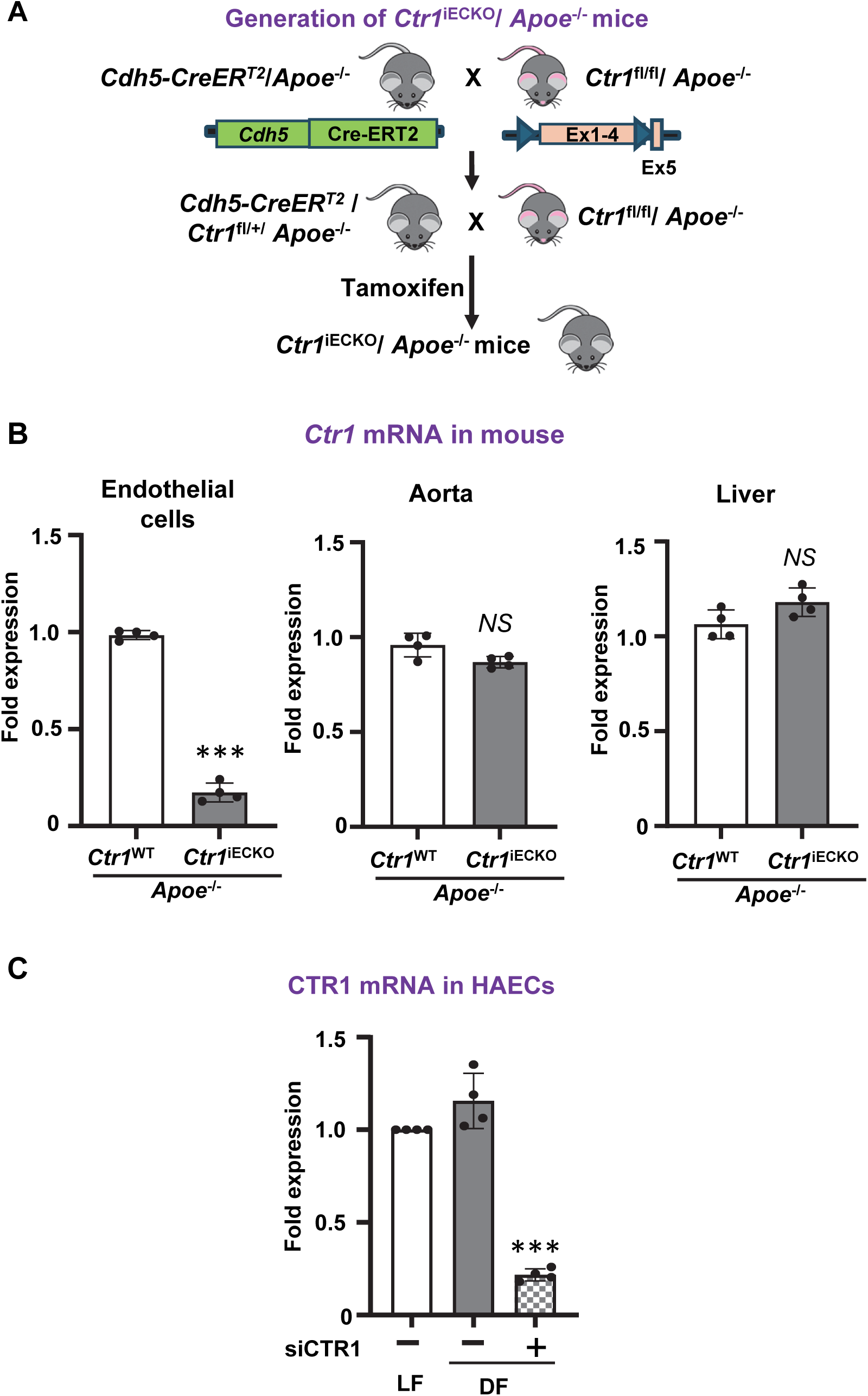
CTR1 expression in endothelial cells and characterization of inducible endothelial *Ctr1*-deficient mice. **A.** Strategy to generate tamoxifen-inducible EC-specific *Ctr1*-deficient (*Ctr1*^iECKO^) mice on *Apoe*^−/−^ background by crossing *Ctr1*^flox/flox^/*Apoe*^−/−^ mice with *Cdh5*-*CreER^T2^*/*Apoe*^−/−^ mice. **B** and **C.** Ctr1 mRNA expression was measured by quantitative polymerase chain reaction in EC or aorta or liver of *Ctr1*^flox/flox^/*Apoe*^−/−^ mice (**B**) and CTR1 siRNA transfected HAEC cells (**C**) n=4.

**Figure S4.**
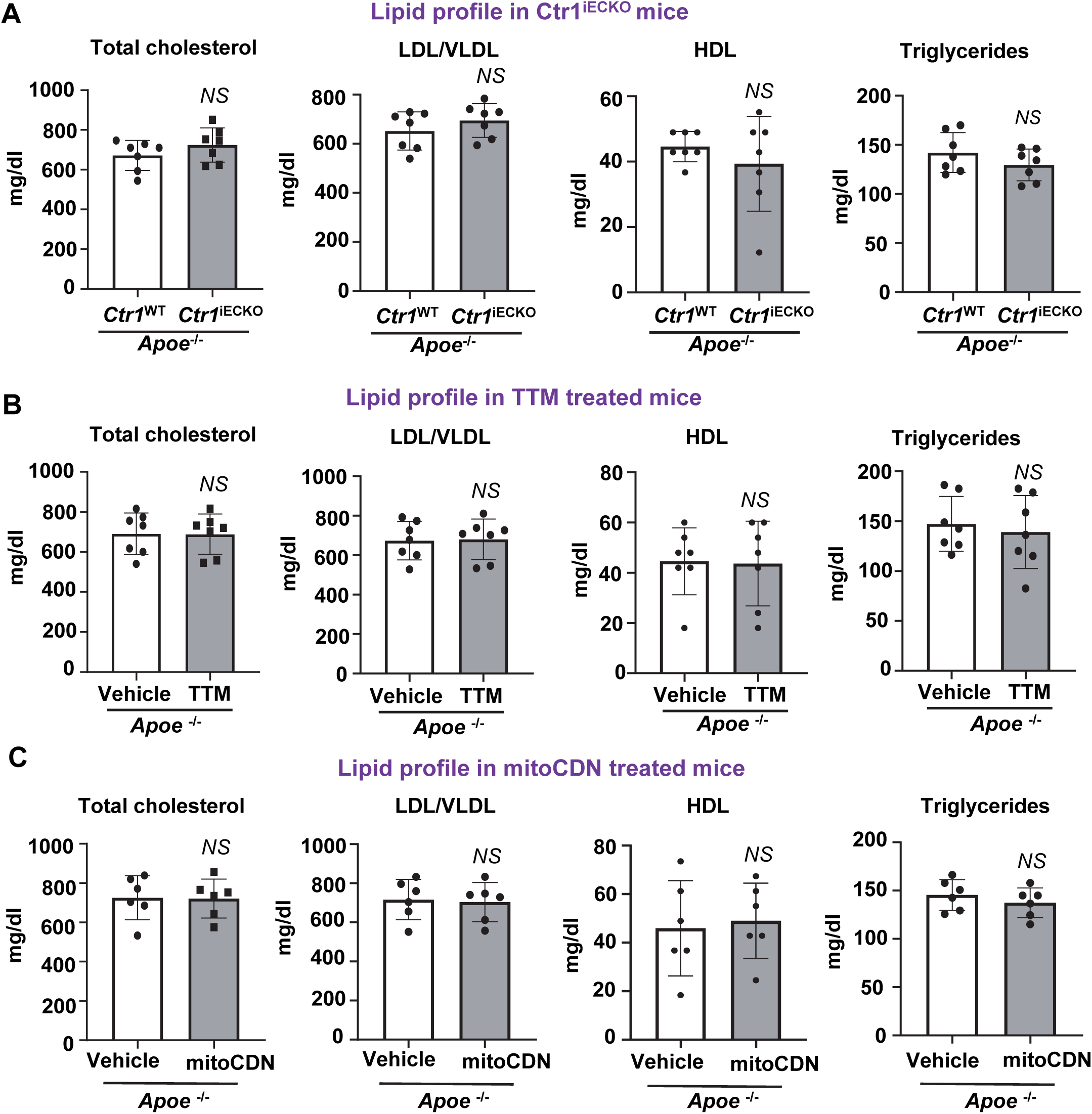
Lipid levels were not changed in TTM treated, EC-Specific *Ctr1*-deficient or mitoCDN-treated *Apoe*^−/−^ mice fed with western diet. Plasma total, LDL or HDL cholesterol and triglyceride levels were measured in mice after overnight fasting in *Ctr1*^iECKO^/*Apoe*^−/−^ (A), TTM treated (B), mitoCDN treated (C) *Apoe*^−/−^ mice (n=6-7).

**Figure S5.**
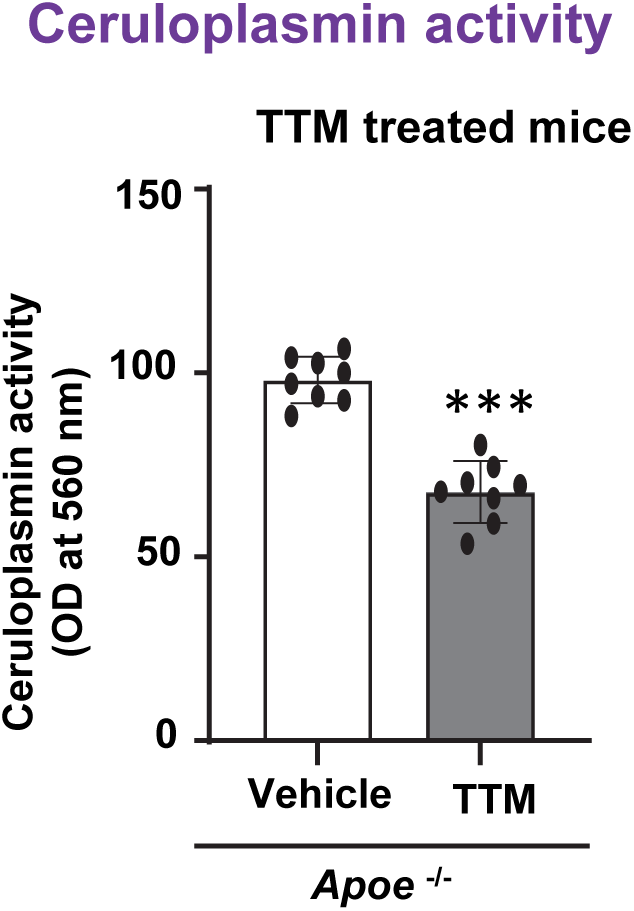
TTM treatment decreased plasma ceruloplasmin activity. Treatment and surgery protocol for these mice are shown in Fig. 1G. **(A)** *Apoe*^−/−^ mice treated with Cu chelator tetrathiomolybdate (TTM) and western diet (WD) received partial carotid ligation (PCL) surgery (3 weeks) on left carotid artery (LCA), n=8. Plasma ceruloplasmin activity was measured.

**Figure S6.**
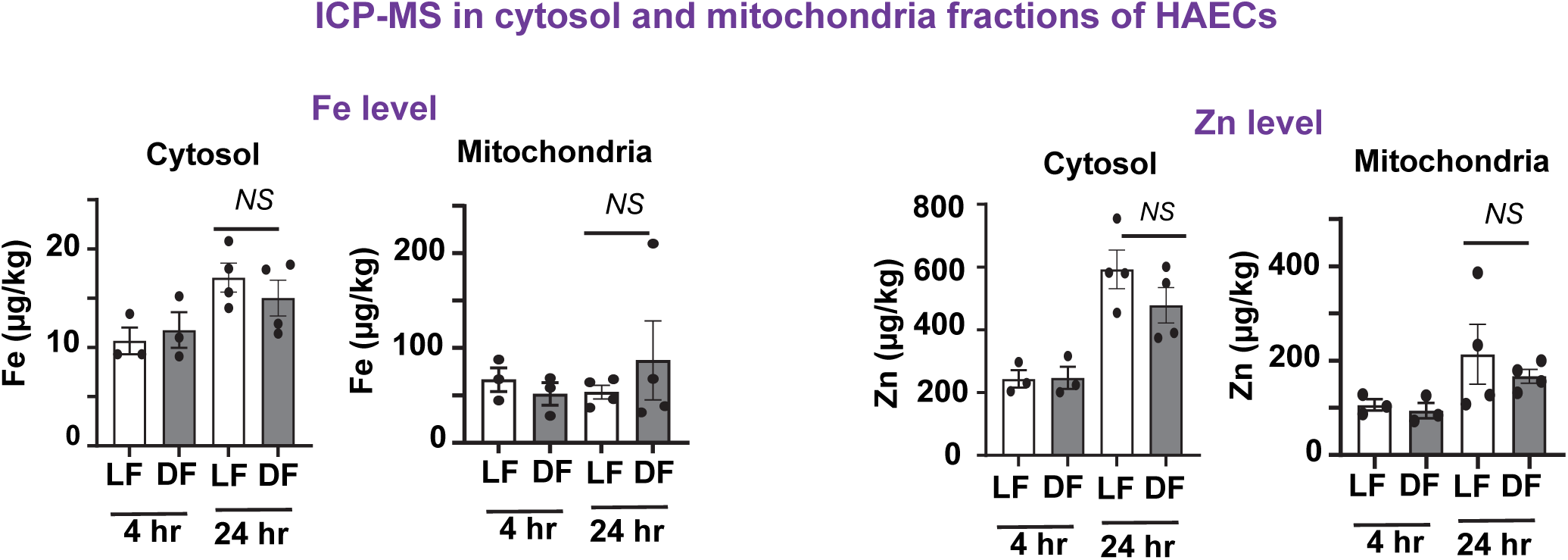
Iron or Zinc contents were not changed in cytosolic or mitochondrial fractions of DF exposed HAECs. After flow exposure in HAECs for 4 or 24 hrs, Fe and Zn levels were measured in cytosolic and mitochondrial fractions of cells (n=3-4) using ICP-MS.

**Figure S7.**
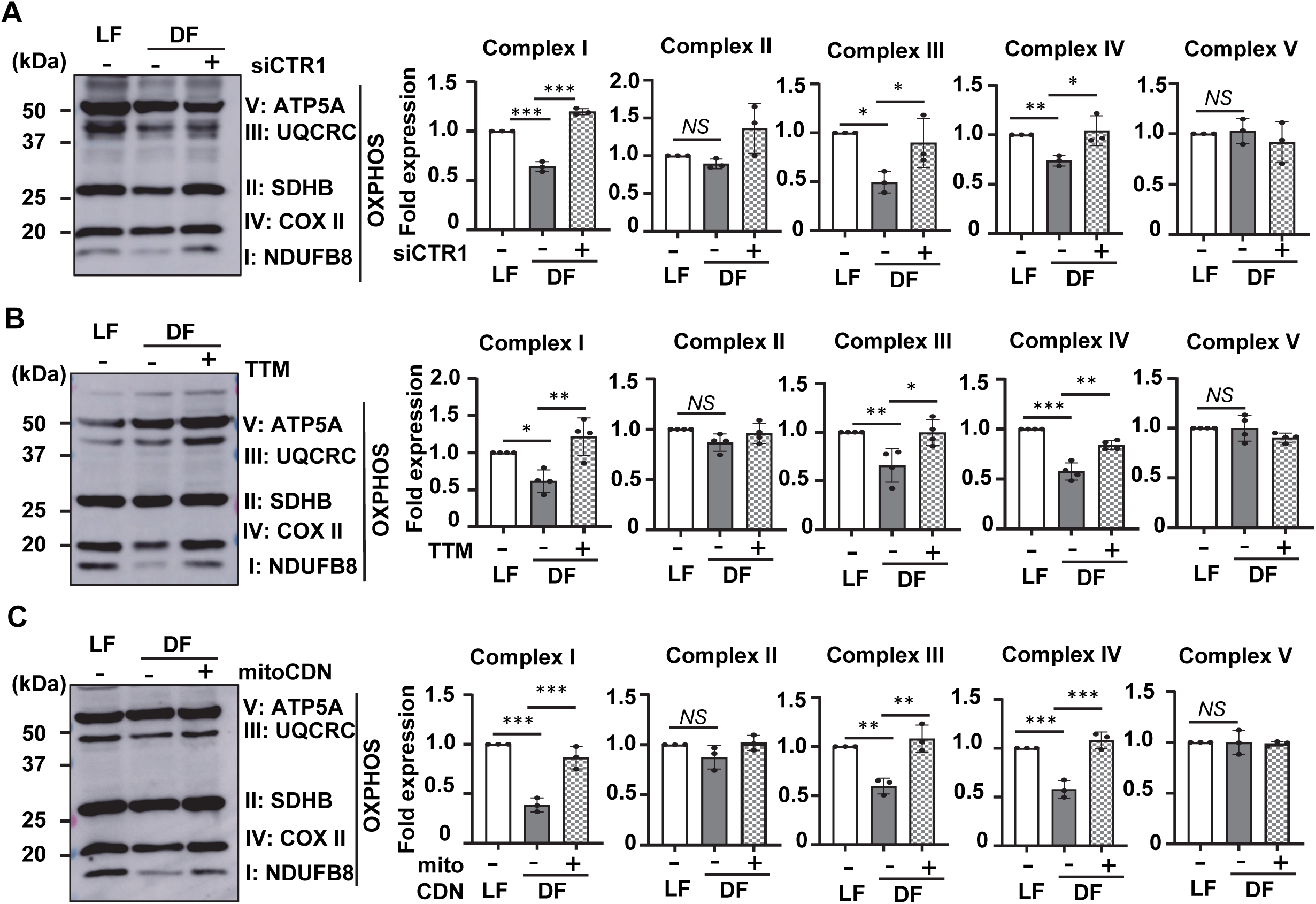
Role of mitochondrial Cu in DF-induced decrease in the protein expression of oxidative phosphorylation complex proteins. **(A-C)** HAEC were transfected with control siRNA or CTR1 siRNA (**A**) or TTM (20nM, 24 hrs) (**B**) or mitoCDN (1 µM, 1 hr) (**C**) and exposed with flow for 48 hrs. Cell lysate were immunoblotted with the indicated antibodies (n=3-4).

**Figure S8.**
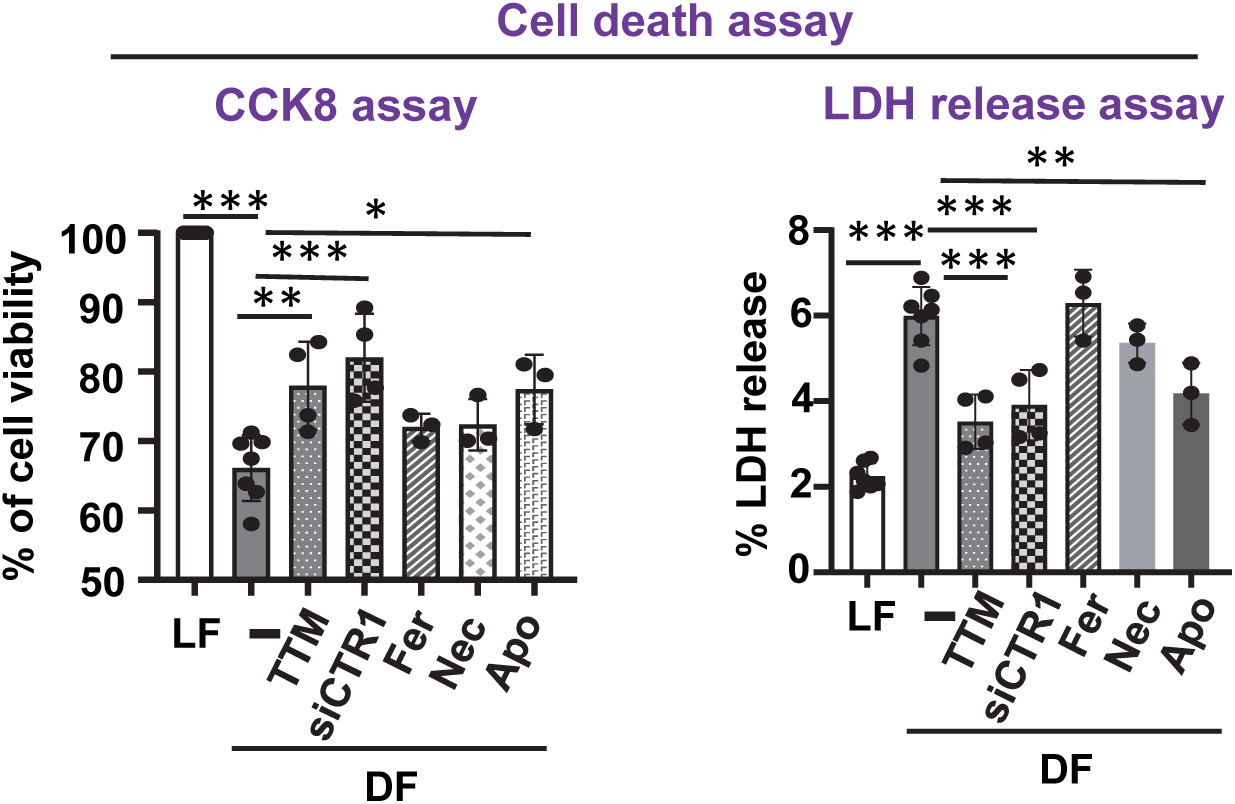
Cu chelator TTM or CTR1 siRNA restored DF-induced decrease in cell viability of ECs. HAEC were transfected with control siRNA or CTR1 siRNA (24 hrs) or pretreated for 1 hr with 20 µM necrostatin-1 (Nec), 10 µM ferrostatin-1 (Fer), 30 µM Z-VAD-FMK (Apo) or pretreated with 20 nM TTM for 24 hrs. The cells were then exposed flow for 48 hrs. Cell death was measured by CCK8 assay (n=4) or LDH release from conditioned medium (n=4).

**Figure S9.**
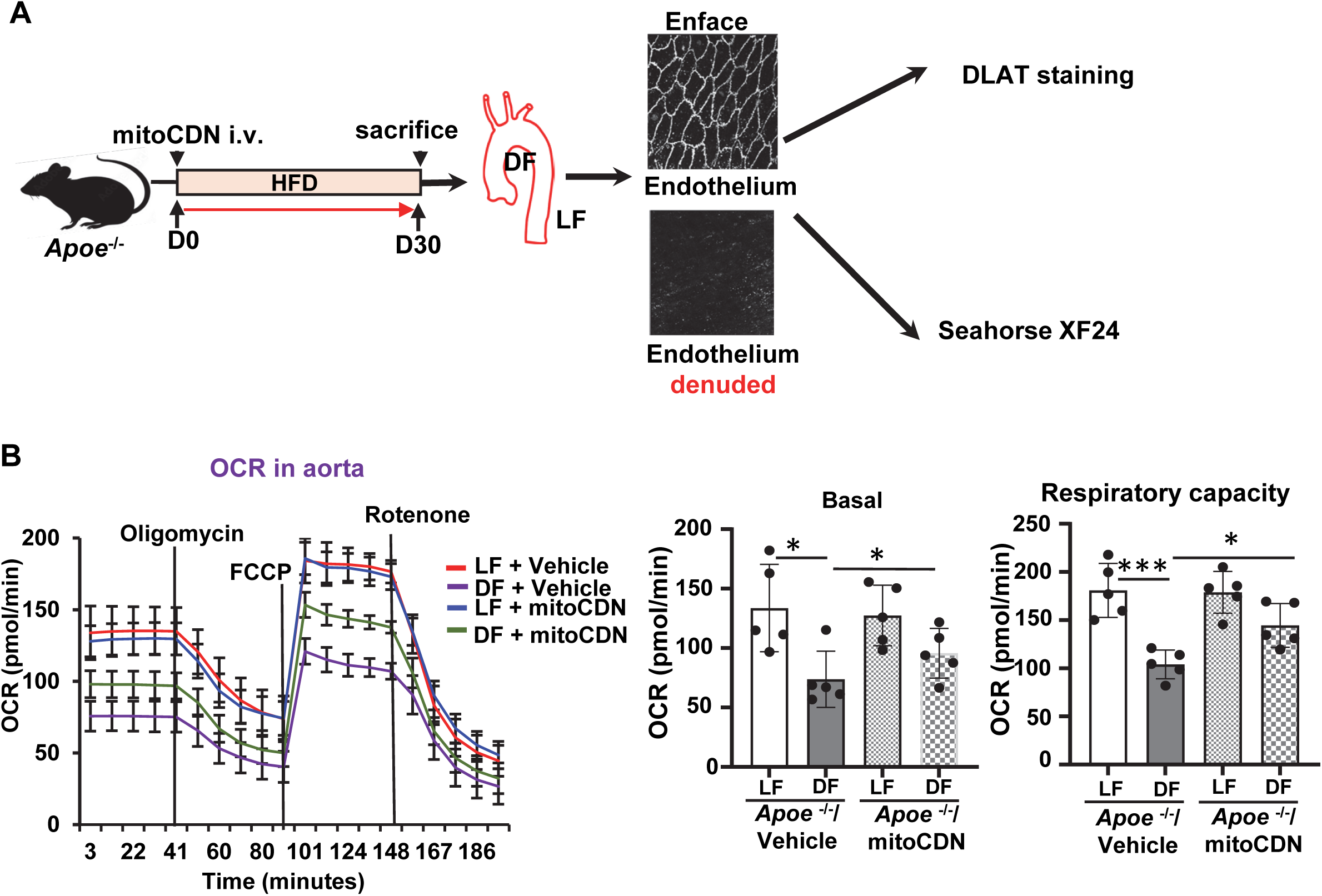
Mitochondrial-specific Cu chelator mitoCDN prevented DF-induced mitochondrial dysfunction *ex vivo*. **A**, Schematic diagram of experimental design for Figure 3G and S9B **B**, *Apoe*^−/−^ mice were treated with mitoCDN (1.35mg/kg) every 3 days with total of 10 doses and fed western diet for 30 days. Mitochondrial respiratory capacity measured in endothelial cells of aortic arch (D-flow, DF) and thoracic aorta (L-flow, LF) of aorta by O**_2_** consumption rate (OCR) using a Seahorse analyzer (n=5).

**Figure S10.**
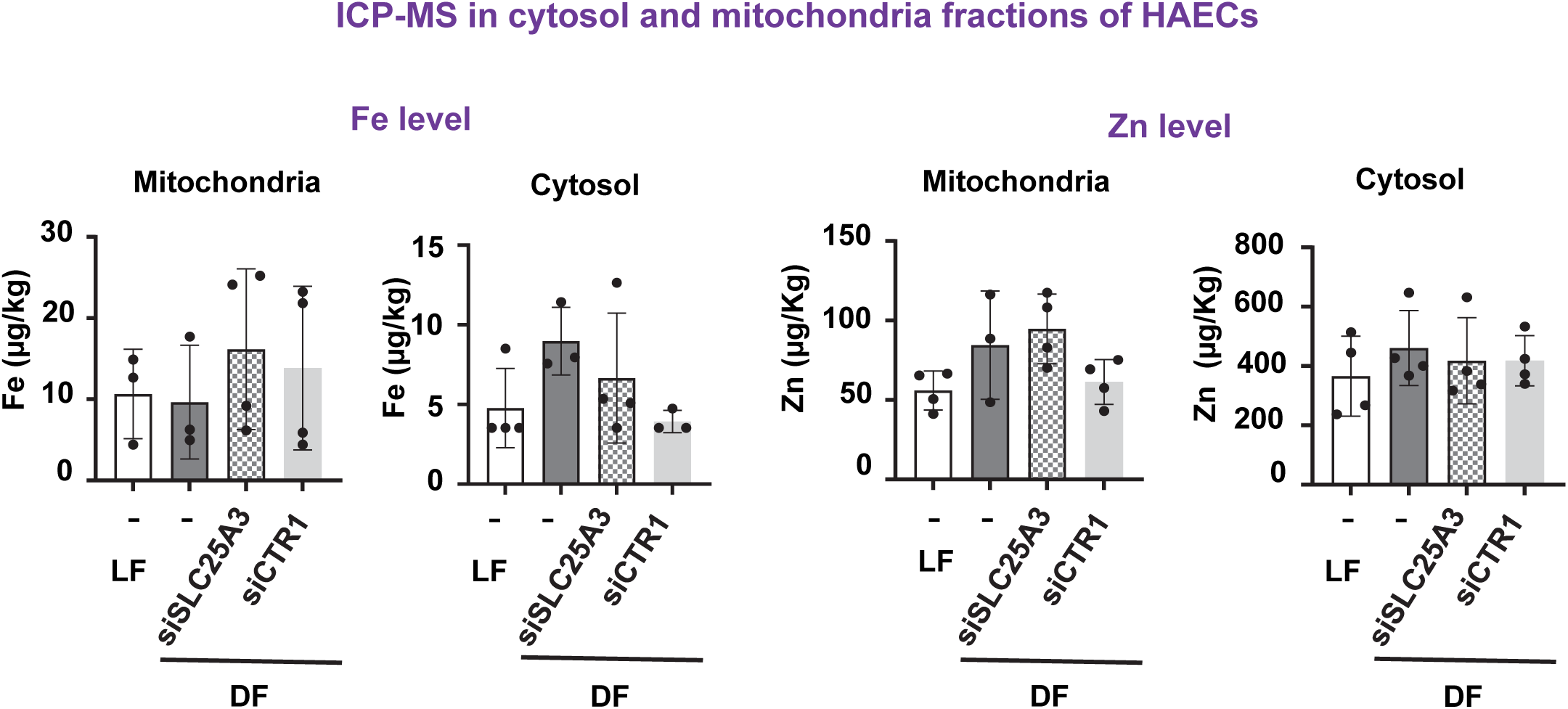
Knockdown of SLC25A3 or CTR1 did not change Fe or Zn levels in flow-exposed ECs. HAECs were transfected with control siRNA or SLC25A3 siRNA or CTR1 siRNA (48 hrs) and exposed to flow for 24 hrs. Iron (Fe) or Zinc (Zn) levels were measured by ICP-MS in cytosolic and mitochondrial fraction of cells (n=3-4).

**Figure S11.**
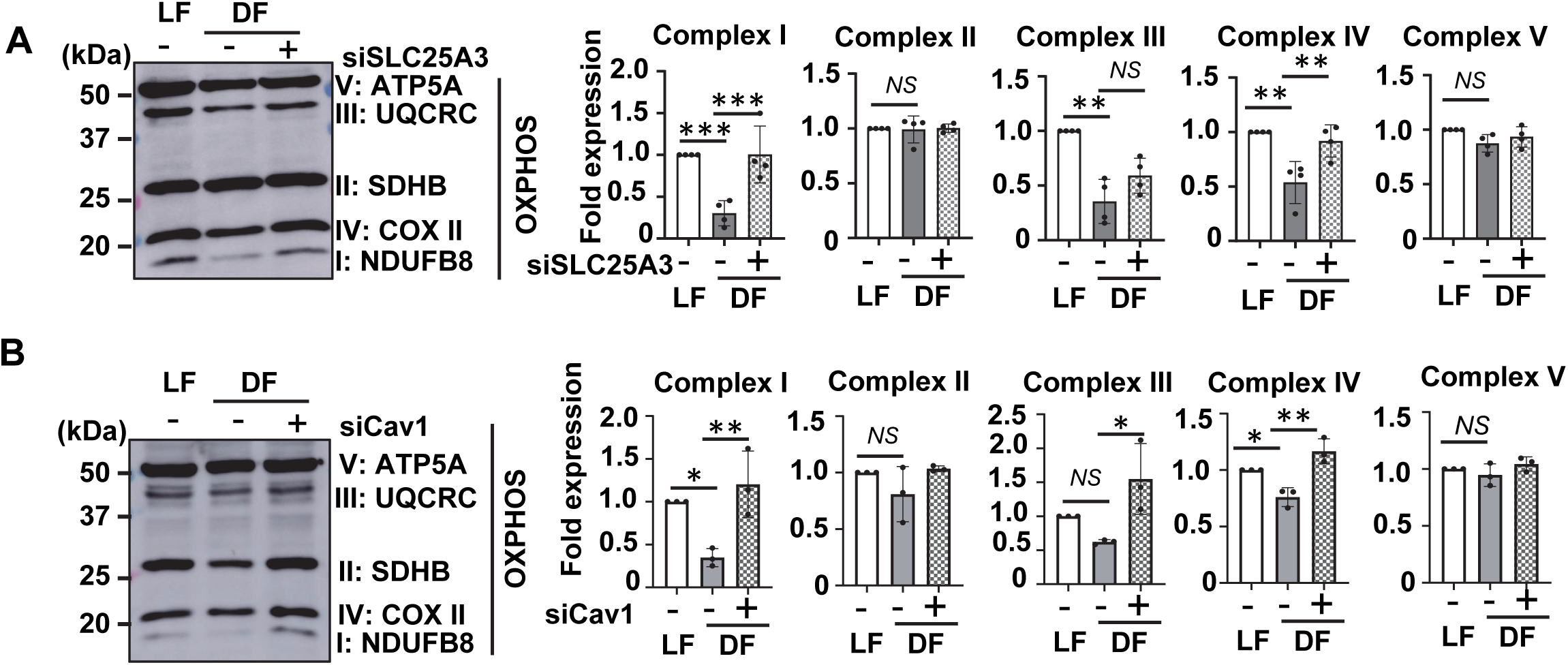
Knockdown of SLC25A3 or Cav1 restored DF-induced decrease in cuproptosis-related protein expression in ECs. (A and. **B)** HAECs were transfected with SLC25A3 siRNA (A) or Cav1 siRNA (B) or control and exposed flow for 48 hrs and cell lysates were immunoblotted with the indicated antibodies (n=3-4).

## Materials and Methods

### Animals

The animal protocols were approved by the by Institutional Animal Care and Use Committee at Medical College of Georgia, Augusta University. And conducted in accordance with federal guidelines and regulations. All mice were maintained at the Augusta University animal facility at controlled temperature (22-23° C, 50-60% humidity) with 12 hrs light/dark cycle and free access to water and standard rodent chow. Mice were held in individually ventilated caging with a maximum of five and a minimum of two mice per cage. Both male and female (8-12 weeks old) mice were used in this study. ApoE-/-mice on the C57BL/6J background were purchased from Jackson Laboratory (Bar Harvor, ME).

### Generation of inducible EC-specific *Ctr1*^−/−^/*Apoe*^−/−^ (*Ctr1*^iEC-KO^/*Apoe*^−/−^) mice

All the mice used in this study were on a C57BL/6J background. First, *Ctr1*^fl/fl^ (*38*) or *Cdh5*-*CreER^T2^* mice were crossed with *Apoe*^−/−^ to produce *Ctr1*^fl/fl^/ *Apoe*^−/−^ or *Cdh5*-*CreER^T2^* / *Apoe*^−/−^ mice, respectively. To generate *Cdh5*-*CreER^T2^* /*Ctr1*^fl/fl^/*Apoe*^−/−^, we crossed *Ctr1*^fl/fl^/ *Apoe*^−/−^ with *Cdh5*-*CreER^T2^* /*Apoe*^−/−^. To induce postnatal deletion of endothelial Ctr1 in adults, we administered tamoxifen (50 mg kg−1 of body weight) by intraperitoneal (i.p.) injection into *Ctr1*^fl/fl^/*Apoe*^−/−^ or *Cdh5*-*CreER^T2^* /*Apoe*^−/−^ (*Ctr1*^WT^) or *Cdh5*-*CreER^T2^* /*Ctr1*^fl/fl^/*Apoe*^−/−^ mice. Injections were performed once per day for 10 days with a 2-day break, followed by a resting period of 2 weeks to obtain Ctr1^iEC-KO^/*Apoe*^−/−^ or *Ctr1*^WT^/*Apoe*^−/−^ mice. We isolated mouse coronary ECs, as described previously(*51*).

### Experimental Design

Partial carotid ligation (PCL) of left carotid artery (LCA) was performed as previously described (*52*). Briefly, LCA was exposed after parenteral anesthesia. Then, the left external carotid, internal carotid and occipital arteries were ligated with 6-0 suture while the superior thyroid artery was left intact. The right carotid artery (RCA) was left unperturbed. The mice were then fed a Western diet (21% fat and 0.2%cholesterol) (ENVIGO, TD88137) for 21 days. Using ultrasonography (VisualSonics VEVO3100 System), development of D-flow in the LCA was confirmed with reversal flow during diastole. After 21 days of PCL surgery, mice were euthanized and the chest and abdominal cavities were opened, and blood was drawn from the right ventricle. Mice were perfused with cold PBS and 4% paraformaldehyde (PFA) through the left ventricle. Using a dissection microscope, the perivascular tissue was carefully dissected from the wall of the carotid artery.

The carotid artery was embedded in optimum cutting temperature compound (Electron Microscopy Sciences, 15712-S) and cryosectioned into 7-μm-thick sections. Sections were mounted onto Trubond 380 Slides (Tru Scientific) and stained with hematoxylin and eosin (H&E) or 2% Oil Red O (Sigma-Aldrich, O0625) to evaluate the size of necrotic cores or atherosclerotic lesions. Slides were stained with anti-Mac3 to evaluate the macrophages, followed by biotinylated anti-rat IgG (Vector Laboratories, BA-4000). The sections were viewed with a Keyence BZ-X710 microscope (Keyence corporation, Itasca, IL, USA). The atherosclerotic lesion and Mac3 were quantified with BZ-X analyzer (Keyence corporation, Itasca, IL, USA).

For TTM treatment, mice were randomly assigned and gavage with water (control) or 0.7 mg/d per 30 g mice TTM performed daily(*53*) for 2 weeks and continue for 21 days after PCL surgery Ceruloplasmin activity was measured in the serum of mice before and after TTM treatment using a colorimetric assay based on substrate oxidation (Sigma-Aldrich, MAK177). The mitoCDN or vehicle were generated, as previously described (*33*). For mitoCDN treatment, mice were randomly assigned intravenous (retro-orbital) injection with vehicle (tris[(2-pyridyl)-methylamine (TPA), 1.35 mg/Kg BW) or mitoCDN (1.35 mg/Kg BW of CDM) for every three days with a total of 8 doses.

### *En face* staining

Enface staining was performed as described previously (*54*). Briefly, the chest and abdominal cavities were opened after euthanization with CO_2_. Aortas were perfused with cold PBS through the left ventricle, followed by 4%PFA. Using a dissection microscope, the periadventitial tissue was carefully dissected from the wall of the aorta and cut open longitudinally and permeabilized with PBS containing 0.1% Triton X-100 and blocked by TBS containing 10% goat serum and 2.5% Tween 20 for 30 min. Aorta were triple stained with anti-VE-Cadherin (Invitrogen, Alexa-fluor 488, A11008), anti-DLAT (Invitrogen, Alexa Fluor 546, A11003) or DAPI (blue). Images were captured by confocal microscopy (Zeiss). The DLAT aggregation was counted.

### Synchrotron X-ray Fluorescence Microscopy

Sections were prepared as described previously (*25*). Sections (5 µm thick) of formalin-fixed paraffin-embedded greater curvature or lesser curvature of aortic arch were used. For X-ray imaging, the sections were mounted intact onto silicon nitride windows (membrane area, 2 × 2 mm; thickness, 200 nm) manufactured by Silson (Blisworth, UK). Specimens were imaged with the scanning X-ray fluorescence microprobe at beamline 2-ID-E of the Advanced Photon Source (Lemont, IL). Undulator-generated x-rays of 10-keV incident energy were monochromatized with a single bounce Si <111> monochromator and focused to a measured spot size of 0.3 x 0.5 µm using Fresnel zone plate optics (X-radia, Concord, CA). Sections were raster-scanned in steps of 4.0 µm, and fluorescence spectra were collected for 1-to 2-sec dwell times by using a 4-element silicon drift detector (Vortex-ME4, SII Nanotechnology, CA). Quantitation and image-processing of the X-ray fluorescence (XRF) data sets was performed with MAPS software (*55*). Quantitation of elemental content was achieved by fitting XRF spectra at each pixel and comparing against a calibration curve derived from measurements of thin-film standard AXO (AXO DRESDEN GmbH, Dresden, Germany).

### Human atherosclerotic arteries

Human atherosclerotic arteries were obtained from cadavers by Dr. Joseph White (Department of Pathology, Augusta University), as described previously (*41*). The heart and aortic tissue were isolated within 24 hrs of death. The isolated tissues were thoroughly rinsed with ice-cold phosphate-buffered saline (PBS), cleaned of perivascular fat and fixed in 2% Paraformaldehyde in PBS, pH 7.4). Aorta embedded in optimum cutting temperature compound (EMS, 15712-S) and cryosectioned into 7-μm-thick sections. Sections were mounted onto Trubond 380 Slides (Tru Scientific). The use of human cadaveric specimens was approved by the Institutional Biosafety Committee at Augusta University (BSP# 1458).

### Mitochondrial respiratory capacity (O_2_ consumption rate, OCR) in aortic arch and thoracic aorta

Mitochondrial stress in vessels was measured as described previously(*56*). Briefly, Aortic arch and thoracic aortas from mice were isolated and kept in seahorse XF Media (Alilent, 103680) supplemented with 5.5 mM glucose and 1 mM pyruvate. The periadventitial tissue was carefully dissected from the wall of the vessel and cut open longitudinally and cut into 2 mm long explants. A portion of the aorta was denuded using cotton swabs. Aortic explants were carefully added to an individual well of Seahorse XF24 Islet Capture Microplates (Agilent, 101122-100). Explants were secured in wells while maintaining their orientation as endothelial cell “side-up” using islet capture screens to measure OCR. After 1h incubation at 37 °C without CO2, the OCR was measured using Seahorse analyzer. Data were analyzed using Wave software (Agilent, Santa Clara, CA).

### Measurement of plasma cholesterol and triglyceride

Mice were fasted overnight before blood was collected. Total cholesterol, LDL, HDL (BioAssay Systems, EHDL-100) and triglyceride (Wako Diagnostics, 992-0292;998-0292) were measured using kit with manufacturer instruction.

### Flow cytometry (FACS) analysis

LCA and RCA were excised after 5 days of PLA surgery and mitoCDN injected every three days with a total of 2 doses (one day prior to surgery and at Day 2 of post-surgery). Carotid arteries collected and pooled (n=3) in collagenase digestion buffer and digested as described (*57*). Briefly, carotid arteries were minced and enzymatically digested in DPBS containing collagenase type I (1 mg/mL) (Sigma, C0130), 18 mg/mL collagenase Type XI (Sigma, C7657), 1 mg/mL hyaluronidase (Sigma, H3506) and 50 U/mL DNAse (Sigma, D4263) at 37°C for 1 hr. Cells were spun and re-suspended in PBS with 2% FCS and filtered using 70 µm and 40 µm filter followed by cell counting. The cell suspension was incubated in blocking buffer consisting of anti-mouse CD16/CD32 (eBioscience, 14-0161-85) in 2% FCS for 20 min in ice. Cells were incubated with APC-labeled anti-mouse CD31 antibody (clone MEC13.3, Biolegend, 102509) and fixable viability dye (Invitrogen, 65-0866-14), at a dilution of 1:100 and 1:1000 respectively in 4°C for 30 min. Unstained cells, single color control and compensation beads (Invitrogen, 01-3333-42) were used for auto-compensation. After staining, cells were washed and fixed with 4% PFA and acquired using ThermoFisher Attune Nxt flow cytometer (Thermofisher Scientific, A24858). The recorded data was analyzed using FlowJO 10.10 software (BD Bioscience).

### Cell Culture and Shear stress experiments *in vitro*

Pooled human aortic endothelial cells (HAECs) were purchased and grown in endothelial growth medium (Cell Application) supplemented with penicillin (100 U/ml), Streptomycin (100 mg/ml) and 10% Fetal bovine serum (FBS) and used for experiments until passage 9.

Cone and plate viscometer(*36*) and parallel plate fluid shear system (*36, 58*) (Flexcell international Corp., Burlington, NC, USA) were used. The parallel plate system was used exclusively for immunofluorescence analysis and the cone and plate viscometer system was used for most of the experiments. L-flow and D-flow were generated through unidirectional flow at 15 dyne/cm^2^ and bidirectionally at ±5 dyne/cm^2^, respectively. HAECs grown on gelatin coated 100-mm tissue culture dishes (Falcon, 353003) for cone and plate viscometer and culture slips microscope slides (Flex cell International Corp., FlexFlow™ Culture Slips) for parallel plate. The shear stress system was kept at 37 °C temperature and 95% humidified air with 5% CO_2_. Cells were further processed for immunofluorescence, Cu measurement, subcellular fractionation, western blot and RNA extraction analysis.

### siRNA and plasmid transfection

**s**iRNAs were obtained from Santa Cruz or Origene. HASM were seeded into culture dishes one day prior to transfection. HASM were grown to 40% confluence and transfected with 30 nM siRNA using Oligofectamine (Invitrogen, 12252011) and Opti-MEM Reduced Serum Medium (Invitrogen, 31985070) for 4 hrs according to manufacturer protocol. For plasmid DNA transfection, cells were transfected with flag-hCTR1-WT (∼4 μg for 100 mm dishes) using polyethylenimine (PEI, Polysciences). After transfection, cells were transferred into growth medium and incubated for 48 h at 37 °C before experiments.

### Cell death assay

HAECs were grown on 100 mm culture dishes and exposed to flow for 48 hrs. Cell death was measured using CCK8 assay and LDH release assay, as follows. For CCK8 assay, Cell Counting Kit 8 (WST-8 / CCK8) (Abcam, ab228554) was used. CCK-8 solution (CCK8: medium, 1:10) was added into each dish and incubated in the dark at 37 °C for 1 hr. The spectrometric absorbance at 450nm was measured using the microplate reader (BioTek). Data were normalized to LF, and the results are presented as the percentage over the LF sample. For LDH activity assay, LDH Cytotoxicity Assay Kit (Cayman chemicals, 601170) was used to measure percentage cytotoxicity in conditioned culture medium, according to the manufacturer’s instructions. The spectrometric absorbance at 490nm was measured using the microplate reader (BioTek). The results are presented as the percentage of LDH release.

### Detergent-free purification of caveolae/lipid rafts (C/LR) membrane fractions

C/LR fractions were separated using the sodium carbonate-based detergent free method (1). Briefly, L-flow and D-flow exposed cells (pooled of 12 x 100 mm dishes) were homogenized in a solution containing 0.5 M sodium carbonate (pH 11), 1 mM sodium orthovanadate, and protease inhibitors. Homogenization was carried out sequentially in the following order using a loose-fitting Dounce homogenizer (10 strokes) and a sonicator (four 20-s bursts). The homogenates were adjusted to 45% sucrose by adding 90% sucrose in a buffer containing 25 mM Mes (pH 6.5) and 0.15 M NaCl and placed at the bottom of an ultracentrifugation tube. A 5–35% discontinuous sucrose gradient was formed above and centrifuged at 39,000 rpm at 4°C for 16–20 h in a Beckman SW-40Ti rotor. From the top of the tube, 13 fractions were collected, and an equal volume from each fraction was subjected to immunoblotting.

### Mitochondria and cytosol fractions

Mitochondria and cytosol fractions were isolated by Mitochondria/Cytosol Fractionation Kit (Abcam, ab65320). Briefly, the cells were washed with ice-cold PBS and suspended in Cytosolic Extraction Buffer containing DTT and protease inhibitors. The cells were incubated on ice for 10 min, and then homogenized on ice with 40 passes using a dounce homogenizer. Homogenates were centrifuged at 700 g for 10 min and the supernatant was collected. The collected supernatant was centrifuged at 10,000 g for 30 min at 4 °C. The supernatant is the cytosolic fraction. The pellets containing the crude mitochondria were lysed in the Mitochondrial Extraction Buffer Mix, and mitochondrial fraction was collected.

### Cu Measurements using ICP-MS

Cells were exposed to D-flow or L-flow, rinsed with PBS and collected into 15 ml metal-free centrifuge tubes (Gibco, 14190250). Then, centrifuged and the cell pellet were submitted. Cytosolic and mitochondrial fractions were also submitted using metal free centrifuge tubes. Pre-digested samples were submitted for ICP-MS analysis to Oregon Health and Science University (OHSU) Elemental Analysis Core. Samples were kept frozen at −80°C until preparation. All samples were further diluted to the 1% HNO3 (trace metal grade, Thermo Fisher Scientific). Inductively coupled plasma mass spectroscopy (ICP-MS) analysis was performed using an Agilent 8900 triple quad equipped with an SPS autosampler. The system was operated at a radio frequency power of 1550 W, an argon plasma gas flow rate of 15 L/min, Ar carrier gas flow rate of 0.9 L/min. Elements were measured in kinetic energy discrimination mode (Fe, Cu, and Zn) using He gas (at 5 ml/min) or in mass-on-mass using O2 (Fe). Data were quantified using weighed, serial dilutions of a multi-element standard (CEM 2, (VHG Labs, VHG-SM70B-100) Fe, Cu, Zn). For each sample, data were acquired in triplicates and averaged. A coefficient of variance (CoV) was determined from frequent measurements of a sample containing ∼10 ppb of Fe, Cu, and Zn. An internal standard (Sc, Ge, Bi) continuously introduced with the sample was used to correct for detector fluctuations and to monitor plasma stability.

### Immunofluorescence analysis

HASM on culture slips microscope slides were rinsed quickly in ice-cold PBS, fixed in freshly prepared 4% paraformaldehyde (Electron Microscopy Sciences, 15712-S) in PBS for 10 min at room temperature, permeabilized in 0.05% Triton X-100 (MP Biomedicals, 194854) in PBS for 5 min, and rinsed sequentially in PBS, 50 μmol/L NH4Cl (Sigma, A4514) and PBS for 10 min each. After incubation for 1 h in blocking buffer (PBS+3%BSA), cells were incubated with anti-flag (Sigma, F3165) or anti-DLAT (Cell signaling, 12362S) or anti-SLC25A3 (Proteintech, 10420-1-AP) for 18 h at 4°C, rinsed in PBS/BSA, and then incubated in Alexa Fluor conjugated goat anti-rabbit IgG (Invitrogen) or Alexa Fluor conjugated goat anti-mouse IgG (Invitrogen) for 1 h at room temperature and cells rinsed with PBS. Cells on coverslips were mounted onto glass slides using Vectashield (Vector Laboratories, H-1200) and observed using confocal microscopy (Zeiss LSM710). The Mito-DsRed vector with the mitochondrial targeting sequence (Invitrogen) was used to visualize and analyze mitochondrial structure as described in manufacturer protocol. Images were taken by confocal microscopy (Zeiss LSM710).

### Intracellular Cu measurement using CF4 probe

Intracellular Cu (+) levels were measured using the copper probe, CF4 (*37*). Cells were exposed to L-flow or D-flow for indicated time in the parallel plate fluid shear system. After cell exposure to shear stress, CF4 (2 µM) was added on the top of slides and incubate for the 5 min. Then, cells were washed twice with ice-cold HBSS buffer and equilibrated for 20 min with HBSS buffer. CF4 fluorescence was imaged by confocal microscopy (Zeiss LSM710). Cells were excited at 594 nm and emission was collected between 610-700 nm.

### Immunoprecipitation and Immunoblotting

Cells were lysed with 500 μl of ice-cold lysis buffer, pH 7.4 (50 mM HEPES, 5 mM EDTA, 120 mM NaCl), 1% Triton X-100, 60 mM n-Octyl-α-D-glucopyranoside, protease inhibitors (10 μg/ml aprotinin, 1 mM phenylmethylsulfonyl fluoride, 10 μg/ml leupeptin) and phosphatase inhibitors (50 mM sodium fluoride, 1 mM sodium orthovanadate, 10 mM sodium pyrophosphate). For immunoprecipitation, cell lysates (700-1000 μg) were precipitated with lipoic acid (Sigma Alrcich, 437695) or CTR1 (Home made) antibody overnight at 4°C and then incubated with 20 μl of protein A/G-agarose beads (Santa Cruz, SC-2003) for 1.5 h at 4°C. Cell lysates (25 μg) or immunoprecipitates were separated using SDS-polyacrylamide gel electrophoresis and transferred to nitrocellulose membranes (Bio-Rad, 1620115), blocked overnight in PBS containing 5% nonfat dry milk and 0.1% Tween 20 (Fisher, BP337), and incubated overnight with primary antibodies. Following primary antibodies were used: Anti-LIAS (Proteintech, 11577-1-AP), Anti-POLD1 (abcam, ab186407), Anti-DLAT (Cell signaling, 12362S), total OXPHOS cocktail (abcam, ab110411), Anti-SLC25A3 (Proteintech, 10420-1-AP), Anti-CTR1 Ab (Abcam, ab129067), Anti-Cav1 (BD Biosciences, 610407), Anti-Paxillin (BD Bioscience, 610051), Anti-VDAC1 (Cell signaling, 4661). After incubation with secondary antibodies (Goat Anti-Rabbit IgG-HRP Conjugate, Bio-Rad#1706515, Goat Anti-Mouse IgG-HRP Conjugate, Bio-Rad#1706516, proteins were detected by ECL chemiluminescence. To check DLAT oligomer detection, we used non-reducing gel.

### Quantitative real-time PCR

Total RNA from HAECs or tissues was isolated by using phenol/chloroform and isolated using Tri Reagent (Molecular Research Center Inc.). Reverse transcription was carried out using high-capacity cDNA reverse transcription kit (Applied biosystems, 4368814) using 2 ug of total RNA. Quantitative PCR was performed with the ABI Prism 7000, the SYBR Green PCR kit (Qiagen, 204054) and the QuantiTect Primer Assay (Qiagen) for specific genes. Samples were run in triplicate. Expression of genes were normalized and expressed as fold changes relative to GAPDH or HRT.

### Mitochondria respiratory capacity (O_2_ consumption rate, OCR) in HAEC cells

Following shear stress (for 48 hrs) exposed cells were trypsinised, plated (250K cells/well, 24 well plate) and seeded in the Seahorse XF24 extracellular flux assay kits (Agilent, 100850-001). After plating for 2 hrs, cell culture medium was changed with modified XF assay medium (100965-000, unbuffered, glucose free, and pyruvate free) containing low glucose (0.1% (W/V), ∼5.5 mM). Cells were incubated at 37 °C without CO2 for 1 hr, the OCR was measured using Seahorse according to manufacturer’s instruction (XF Cell Mito stress test kit;103015-100).

### Statistical Analysis

Data are presented as mean ± SEM. The statistical analysis was based on sample size (n), indicating the number of biologically independent experiments, biologically independent samples, biologically independent animals, individual cells or fields of view as described in detail in the respective figure legends. No statistical method was used to predetermine the sample size. Each experiment was performed a minimum of three times to ensure reproducibility. No samples or animals were excluded from the analysis. All mice/samples were randomized before experiments. Data collection and analysis were performed blinded to group allocation. Data were compared between groups of cells and animals by Student *t* test when one comparison was performed or by ANOVA for multiple comparisons. When significance was indicated by ANOVA, the Tukey post-hoc and Bonferroni multiple comparison analysis was used to specify between group differences. Values of *p<0.05, **p<0.01 and ***p<0.001 were considered statistically significant. Statistical tests were performed using Graphpad Prism 10 (GraphPad Software, San Diego, CA).

## Author contribution

T.F., M.U.-F. and V.S., designed the study; V.S (major), A.D., D.A., S.Y., B.C., performed/assisted research; M.U.-F., T.F., V.S., S.Y., analyzed data; G.C., provided human atherosclerosis sample; O.A., S.V., performed XFM experiments; C.D.M., A.T.P., and C.J.C. developed and provided the CF4 copper imaging probe; H.J., provided inputs and edited manuscript; J.L., provided Ctr1 fl/fl mice; J.H.K., provided inputs on experimental design and data interpretation, and edited manuscript; M.M. and S.K. performed mouse genotyping; Z.X., and J.R., developed and provided mitoCDN; T.F., M.U.-F. and V.S. wrote the manuscript. All of the authors reviewed the manuscript. XFM experiments were performed using resources of the Advanced Photon Source, a U.S. Department of Energy (DOE) Office of Science user facility operated for the DOE Office of Science by Argonne National Laboratory under Contract No. DE-AC02-06CH11357. We acknowledge the assistance of Dr. Martina Ralle, Director of USR Elemental Analysis Core at OHSU (partial support from NIH (S10OD028492) for measuring metals levels using ICP-MS.

## Data and materials availability

All data necessary to evaluate the conclusions in this paper are present either in the main text or the supplementary materials. Additional data supporting the findings of this study are available from the corresponding author upon reasonable request.

